# Single-cell chromatin profiling reveals demethylation-dependent metabolic vulnerabilities of breast cancer epigenome

**DOI:** 10.1101/2020.02.18.954495

**Authors:** Meena Kusi, Maryam Zand, Chun-Lin Lin, Chiou-Miin Wang, Nicholas D. Lucio, Nameer B. Kirma, Jianhua Ruan, Tim H.-M. Huang, Kohzoh Mitsuya

## Abstract

Metabolic reprogramming in cancer cells not only sustains bioenergetic and biosynthetic needs but also influences transcriptional programs, yet how chromatin regulatory networks are rewired by altered metabolism remains elusive. Here we investigate genome-scale chromatin remodeling in response to 2-hydroxyglutarate (2HG) oncometabolite using single-cell assay for transposase accessible chromatin with sequencing (scATAC-seq). We find that 2HG enantiomers differentially disrupt exquisite control of epigenome integrity by limiting α-ketoglutarate (αKG)-dependent DNA and histone demethylation, while enhanced cell-to-cell variability in the chromatin regulatory landscape is most evident upon exposure to L2HG enantiomer. Despite the highly heterogeneous responses, 2HG largely recapitulates two prominent hallmarks of the breast cancer epigenome, i.e., global loss of 5-hydroxymethylcytosine (5hmC) and promoter hypermethylation, particularly at tumor suppressor genes involved in DNA damage repair and checkpoint control. Single-cell mass cytometry further demonstrates downregulation of BRCA1, MSH2 and MLH1 in 2HG-responsive subpopulations, along with acute reversal of chromatin remodeling upon withdrawal. Collectively, this study provides a molecular basis for metabolism-epigenome coupling and identifies metabolic vulnerabilities imposed on the breast cancer epigenome.

## Introduction

As a dynamic system, cancer cells continuously adapt to the fluctuating microenvironment by rerouting metabolic fluxes and evolve from early initiation through progression and dissemination^1,2^. Metabolic reprogramming in cancer cells facilitates energy production and macromolecular synthesis to fuel cell proliferation^3,4^. In addition to supporting bioenergetic and biosynthetic needs, altered metabolism involves promiscuous production of non-canonical metabolic intermediates, which have been described as metabolic waste products or metabolite damage^5,6^. Recent studies suggest that these previously uncharacterized metabolites including oncometabolites are linked to ‘non-metabolic’ signaling mechanisms in cell-type-specific fate decisions^7^.

The oncometabolite 2-hydroxyglutarate (2HG) occurs as two enantiomers and accumulates up to millimolar concentrations in a broad range of hematological and solid malignancies^8,9^. Somatic mutations in isocitrate dehydrogenase genes, *IDH1* and *IDH2*, found in glioma and acute myeloid leukemia (AML) result in stereospecific production of the D-enantiomer (D2HG)^10,11^, while breast tumors frequently exhibit elevated levels of 2HG despite the lack of *IDH* mutations^12,13^. Recent studies indicate that the L-enantiomer (L2HG) can accumulate under acidic^14,15^ and hypoxic conditions^16,17^ that often coexist in the tumor microenvironment, yet the potential sources and functions of L2HG are less well established. Both enantiomers structurally resemble α-ketoglutarate (αKG), a key intermediate in the Krebs cycle, and potentially antagonize αKG-dependent dioxygenases including ten-eleven translocation (TET) DNA hydroxylases and Jumonji domain-containing histone demethylase (JHDM) enzymes^18–20^, which catalyze oxidative demethylation of DNA and histone proteins, respectively.

Promoter hypermethylation and global loss of 5-hydroxymethylcytosine (5hmC) are two prominent hallmarks of the breast cancer epigenome^21–23^. Genome-wide depletion of 5hmC is frequently observed in a multitude of tumor types including breast cancer and is associated with poor patient survival^24,25^; however, its cause and pathological consequences are largely opaque. Similarly, promoter hypermethylation of tumor suppressor genes has also been considered a common and driving event in breast malignancies. Loss-of-function mutations in DNA hydroxylases (*TET1*, *TET2* and *TET3*) or overexpression of DNA methyltransferases (*DNMT1*, *DNMT3A* and *DNMT3B*)^26,27^, and recently tumor hypoxia^28^ have been reported to be associated with aberrant DNA methylation, yet the molecular origin of promoter hypermethylation in breast cancer remains obscure^27,29^.

Here we show that the oncometabolite 2HG disturbs the fine-tuned spatial regulation of the mammary epithelial epigenome, and initiates global loss of 5hmC and tumor-associated promoter hypermethylation by impairing αKG-dependent demethylation. The findings provide a mechanistic framework in which altered metabolism mediates two pathological hallmarks of the breast cancer epigenome, thereby priming early epigenetic events to be exploited during the development of breast cancer, in particular basal-like subtype with high 2HG accumulation. By leveraging two single-cell approaches, our study further highlights a role for 2HG oncometabolites in the dynamics of cell-to-cell variability in the epigenome of this tumor type, whereby chromatin regulatory modules are highly vulnerable to metabolic derangements.

## Results

### 2HG enantiomers progressively modulate the mammary epigenome by limiting DNA and histone demethylation

To identify the regulatory mechanism involving cellular metabolism that mediates breast cancer development, we first analyzed the intracellular levels of D2HG and L2HG using enantiomer-selective liquid chromatography-mass spectrometry (LC-MS) following chiral derivatization of the analyte (Fig. 1a,b). Relatively higher levels of D2HG were observed in both ERα-positive and - negative breast cancer cell lines compared to benign and primary mammary epithelial cells, whereas elevated L2HG levels were predominant in ERα-negative breast cancer cells (Fig. 1c,d). In line with this, ERα-negative cells had higher levels of total 2HG (Supplementary Fig. 1a), which is supported by prior studies showing preferential accumulation of 2HG in ERα-negative or basal-like tumors^12,13,30^. We next leveraged the recently developed comprehensive Cancer Cell Line Encyclopedia (CCLE) metabolomics database^31,32^ and found that 2HG levels were remarkably elevated in basal-like tumor cells in comparison to other tricarboxylic acid (TCA) cycle metabolites that are structurally similar to one another and are potentially involved in αKG-dependent dioxygenase reactions (Fig. 1e). Of note, our investigation of somatic mutations using two independent datasets from the Sanger Catalogue of Somatic Mutations in Cancer (COSMIC) and CCLE consortia showed no evidence for gain-of-function mutations in *IDH1* or *IDH2* among 62 breast cancer cell lines, further corroborating the findings of infrequent *IDH* mutations and yet high accumulation of 2HG in breast tumors (Supplementary Fig. 1b)^12,13,30,33^. It should be noted, however, that recurrent *IDH2* mutations are found in a rare breast cancer subtype with elevated intratumor 2HG levels^34,35^.

**Fig. 1.**
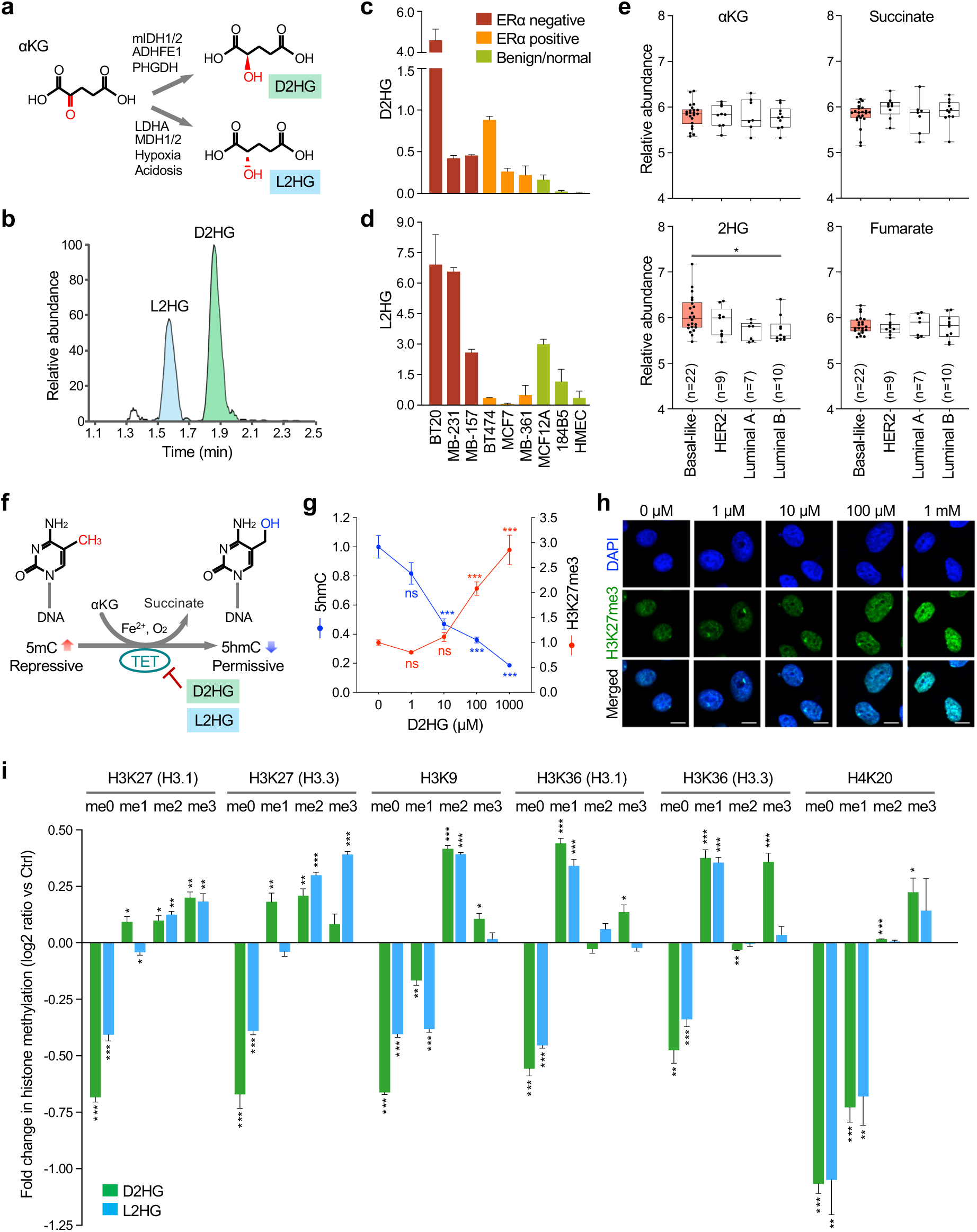
2HG imposes a global loss in 5hmC and progressive gain in histone methylation on the mammary epithelial epigenome. **a**, Schematic of reductive conversion of αKG to the enantiomers of 2HG. 2HG is stereospecifically produced from αKG by mutant IDH proteins (mIDH1/2) or by promiscuous enzymatic reactions. **b**, Extracted-ion chromatogram showing chiral derivatives of 2HG detected in breast cancer cells. **c**,**d**, Chiral LC-MS analysis of the two enantiomers D2HG (**c**) and L2HG (**d**) in breast cancer cell lines, benign and normal mammary epithelial cells (n = 3 independent replicates). **e**, Intratumor levels of αKG, 2HG, succinate and fumarate in the CCLE breast cancer cell lines. αKG, 2HG, succinate and fumarate are structurally similar to one another and are potentially involved in DNA and histone demethylation processes. \**P* < 0.05 versus basal-like tumor cells by one-way ANOVA with Dunnett’s multiple comparison test. **f**, Schematic of 2HG-mediated inhibition of αKG-dependent DNA demethylation by the TET family of DNA hydroxylases. TET enzymes catalyze oxidation of 5mC to 5hmC with oxygen and Fe^2+^ as cofactors. **g**, Global levels of 5hmC and H3K27me3 following 72-hr exposure to D2HG. Line plot is shown for the mean intensity of 5hmC (blue) and H3K27me3 (red) assayed by immunofluorescence staining. \*\*\**P* < 0.001 versus control cells by one-way ANOVA with Dunnett’s multiple comparison test (n = 8 images per condition). ns, not significant. At least 60 nuclei were examined and signal intensities were normalized to DAPI nuclear counterstain. **h**, Representative immunofluorescence images of HMECs after 72-hr exposure to D2HG at the indicated concentrations. Scale bar, 10 μm. **i**, Fold change in different types of histone methylation upon 2HG exposure. Following 72-hr exposure to 100 μM of either D2HG or L2HG, global levels of methylated/unmethylated lysine residues were assayed by multiplexed mass spectrometry. While two major histone H3 variants H3.1 and H3.3 are highly similar in their amino acid composition, H3.1 is predominantly localized at heterochromatin, which is in contrast to the preferential H3.3 accumulation in intragenic regions of transcribed genes. \**P* < 0.05, \*\**P* < 0.01, \*\*\**P* < 0.001 versus control unexposed HMECs by two-tailed unpaired Student’s *t*-test with Holm-Sidak correction for multiple comparisons (n = 3 independent replicates). Error bars in column charts or line plot represent s.e.m. αKG, α-ketoglutarate; 2HG, 2-hydroxyglutarate; IDH, isocitrate dehydrogenase; ER, estrogen receptor; LC-MS, Liquid chromatography-mass spectrometry; CCLE, Cancer Cell Line Encyclopedia; HER2, human epidermal growth factor receptor 2; 5hmC, 5-hydroxymethylcytosine; 5mC, 5-methylcytosine; TET, ten-eleven translocation; me0, unmethylated; me1, monomethylated; me2, dimethylated; me3, trimethylated histone lysine residues.

To determine the global impact of oncometabolites on the chromatin landscape, primary human mammary epithelial cells (HMECs), which exhibited low levels of endogenous 2HG (Supplementary Fig. 1a), were exposed for 72 hr to the cell-permeable derivatives of D2HG and L2HG. Exposure to either of the two enantiomers led to a decrease in 5-hydroxymethylcytosine (5hmC) and reciprocal increase in 5-methylcytosine (5mC) across the LINE-1 repetitive sequence elements in the genome (Supplementary Fig. 1c), indicating competitive inhibition of TET-mediated oxidation of 5mC to 5hmC during the process of active DNA demethylation^36^ (Fig. 1f). Similarly, immunofluorescence staining using a 5hmC-specific antibody showed a dose-dependent global loss of 5hmC in response to D2HG (Fig. 1g). In contrast, global levels of H3K27me3 histone methylation were elevated in proportion to the increasing concentrations of D2HG, via inhibition of JHDM-mediated histone demethylation (Fig. 1g,h and Supplementary Fig. 1d). Our observations thus indicated that short-term exposure at a relatively low range was sufficient to induce a substantial change in the mammary epithelial epigenome.

Unlike DNA methylation, lysine residues on histone tails can be mono-, di- and tri-methylated (Supplementary Fig. 1d) and we therefore employed a multiplexed LC-MS assay to quantify changes in histone methylation. Exposure to 2HG enantiomers led to elevated methylation levels in both repressive (H3K27me3, H3K9me3 and H4K20me3) and permissive (H3K4me3, H3K36me3 and H3K79me3) histone marks as indicated by increases in mono-, di- or tri-methylation along with a reciprocal decrease in the levels of unmethylated lysine residues (Fig. 1i, Supplementary Fig. 1e and Supplementary Table 1). Consistently, repressive methylation modifications on H3K27 residues were accumulated in relation to the increasing levels of intracellular 2HG in the CCLE breast cancer cell lines (Supplementary Fig. 1f). Of note, there was no detectable alteration in cell-cycle phase distribution upon 2HG supplementation (Supplementary Fig. 1g). Together, our findings support a model that the intratumor accumulation of 2HG oncometabolites progressively modulates the mammary epigenome independent of cell cycle progression, potentially leading to extensive chromatin remodeling.

### Single-cell profiling of chromatin accessibility reveals epigenetic heterogeneity in response to 2HG enantiomers

To delineate the genome-scale dynamics of epigenetic regulatory modules in response to oncometabolite exposure, we applied single-cell assay for transposase-accessible chromatin with high-throughput sequencing (scATAC-seq)^37^. First, individual single cells were isolated from HMECs cultured in media supplemented with either D2HG or L2HG using integrated fluidic circuits (Fig. 2a). Following cell lysis and tagmentation of open, accessible chromatin regions by Tn5 transposase, a total of 248 single-cell libraries were independently indexed with unique barcodes and sequenced as a single pool. scATAC-seq reads exhibited the expected periodicity of ~200-bp insert size fragments corresponding to nucleosome bands^37,38^ (Fig. 2b). Additionally, aggregate scATAC-seq profiles of 248 single cells showed high concordance with the ensemble measurement of the accessibility landscape profiled by DNase-seq (Fig. 2c and Supplementary Fig. 2a; *r* = 0.76).

**Fig. 2.**
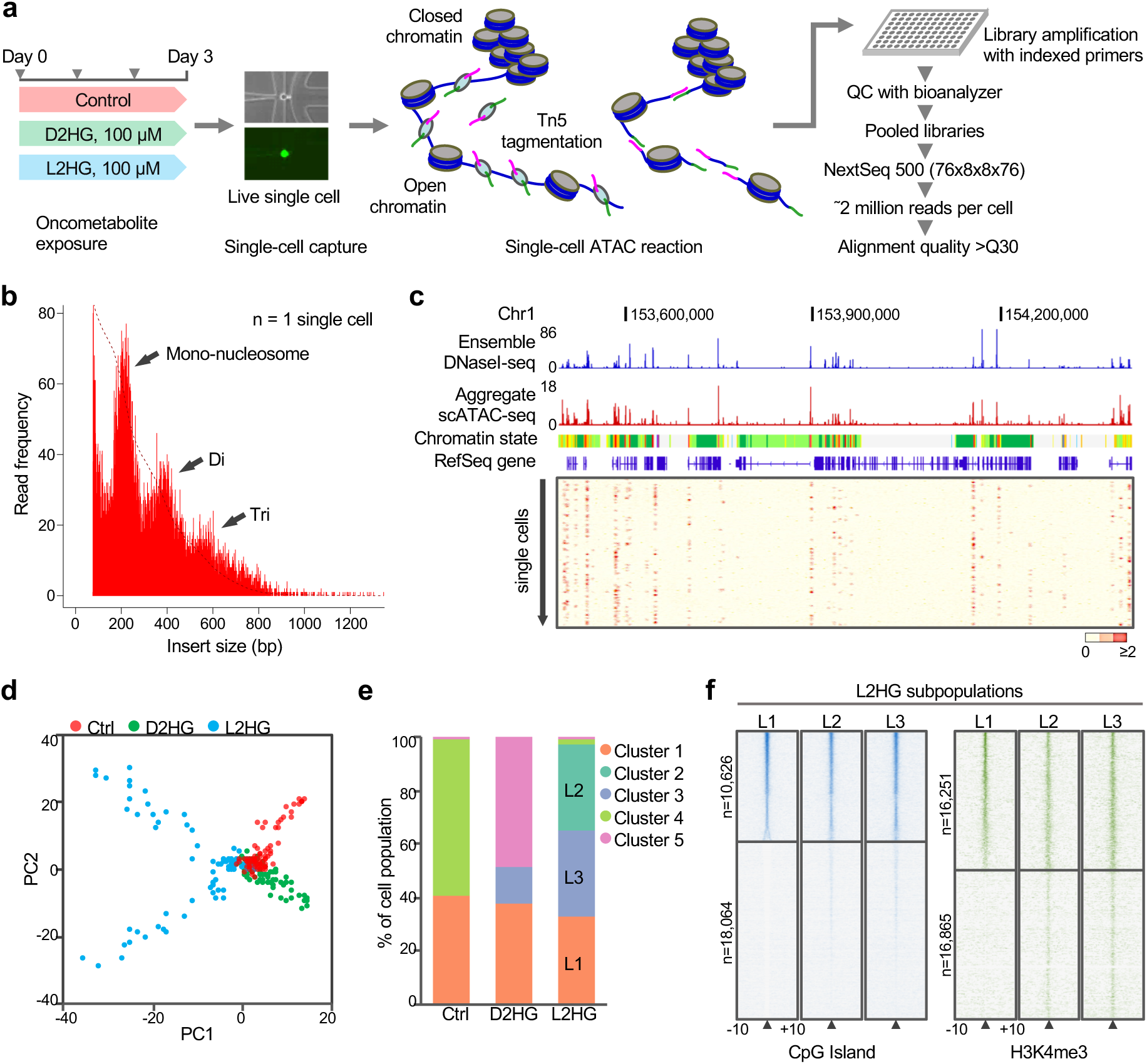
Single-cell ATAC-seq identifies epigenetically heterogeneous responses to 2HG exposure. **a**, Schematic view of the scATAC-seq protocol. Culture medium was replaced daily by freshly prepared media with or without oncometabolite supplementation (arrowheads). **b**, Insert size distribution of ATAC-seq fragments from a single cell displaying characteristic nucleosome-associated periodicity. Diagnostic insert sizes for mononucleosome, dinucleosome and trinucleosome are labeled. **c**, Representative genome browser tracks showing open chromatin regions in HMECs detected by scATAC-seq, which are highly consistent with DNase hypersensitive sites (DHS) detected by bulk DNaseI-seq (GSE29692). Single cell profiles are shown below the aggregated profiles. **d**, Principal component analysis (PCA) of 248 single cells to identify cell subpopulations based on the chromatin accessibility landscape. Ctrl (n = 83), D2HG- (n = 90) and L2HG-exposed (n = 75) cells are colored by sample type. Each data point represents a single cell. **e**, Proportions of cell subpopulations identified by model-based clustering. Cell clusters are color-coded as indicated. **f**, Whole-genome heat maps showing enrichment of scATAC-seq signals in L2HG cell subsets over a 20-kb region centered on CpG islands and H3K4me3 peak summits. Each row represents one individual genomic locus and is sorted by decreasing scATAC-seq signal. The color represents the intensity of chromatin accessibility. Intervals flanking indicated feature are shown in kilobases. ATAC, assay for transposase accessible chromatin; HMECs, human mammary epithelial cells.

We next sought to globally visualize single-cell DNA accessibility profiles using principal component analysis (PCA), which indicated distinct cell subpopulations that were discernible not only between but also within the treatment groups (Fig. 2d and Supplementary Fig. 2b). Cell-to-cell variability in the chromatin landscape was likewise detected among cells receiving the same treatment by using cell similarity matrix analysis based on highly accessible genomic regions (Supplementary Fig. 2c). In contrast, replicate samples were similarly distributed despite being processed in different experiments (Supplementary Fig. 2d), further supporting the technical robustness and reproducibility of the method. Interestingly, the first principal component (PC1) broadly separated L2HG-exposed cells from control and D2HG-exposed cells that were distinguishable, albeit to a lesser extent, alongside the PC2 direction (Fig. 2d), indicating that the two enantiomers could differentially modulate the chromatin organization. Furthermore, L2HG-exposed cells were clearly separated into different subpopulations oriented in the opposite direction on the PC2 axis and showed a more variable distribution compared to unexposed control cells, suggesting a substantial increase in cell-to-cell variability following L2HG exposure.

Consistent with the observation that epigenetic heterogeneity was most evident among L2HG-exposed cells, model-based clustering following PCA identified three major cell subsets in L2HG-exposed cells, referred to as L1, L2 and L3, along with a total of five distinct clusters among the 248 single cells (Fig. 2e and Supplementary Fig. 2e). When scATAC-seq data were aggregated, notable differences in chromatin accessibility were observed throughout the genome between L2HG subgroups (Supplementary Fig. 2f). Moreover, inspection of genome-wide distribution of scATAC-seq signals over CpG islands (n = 28,690) and H3K4me3 peaks (n = 33,116) revealed a marked difference between L1 versus L2 and L3 subsets (Fig. 2f). In agreement, Cluster 1 comprised cells from all three experimental groups that were clustered in close proximity with similar multidimensional phenotypes (Fig. 2d,e), implying L1 subset being a cell subpopulation that is potentially less sensitive to oncometabolite perturbation. This was contrasted with no pronounced differences in scATAC-seq accessibility profiles either across CpG islands or H3K4me3 sites detected between the three experimental groups (Supplementary Fig. 2g). Together, single-cell DNA accessibility profiling revealed that the two enantiomers could distinctly modulate the mammary epigenome and drive cell-to-cell epigenetic diversity in the chromatin regulatory landscape.

### 2HG depletes promoter accessibility at highly methylated genes in breast cancer

To identify disease-relevant epigenetic signatures imposed upon 2HG exposure, we further interrogated our scATAC-seq data using publicly available large-scale datasets including Encyclopedia of DNA Elements (ENCODE)^39^, Roadmap Epigenomics^40^, Cancer methylome^41^ and The Cancer Genome Atlas (TCGA)^42^ (Fig. 3a). We first determined the global occupancy of open, accessible chromatin regions by utilizing the 12-state chromatin segmentation defined in HMECs^43^. As expected, ATAC-seq signals in unexposed control cells were remarkably enriched in active/weak gene promoters and strong enhancers but were significantly underrepresented in inactive genomic regions such as heterochromatin and repetitive regions, both of which were associated with nuclear lamina^40^ (Supplementary Fig. 3a). Following 2HG exposure, chromatin accessibility was reduced in genomic regions enriched with permissive chromatin marks (H3K4me3, H3K27ac and H3K9ac)^40^, such as active/weak promoters and strong/weak enhancers (Fig. 3b). In contrast, genomic regions that are largely devoid of permissive chromatin marks displayed a substantial increase in chromatin accessibility, suggesting that oncometabolites can have two opposing impacts on the mammary epigenome, i.e., selective loss of accessibility in active or poised chromatin and gain of accessibility in repressive or quiescent chromatin states. The overall alterations in the chromatin landscape dynamics were particularly prominent in response to L2HG, suggesting L2HG to be more potent in modulating the mammary epigenome. This is in line with earlier biochemical studies indicating that L2HG can competitively antagonize αKG-dependent dioxygenases to a greater extent^18–20^.

**Fig. 3.**
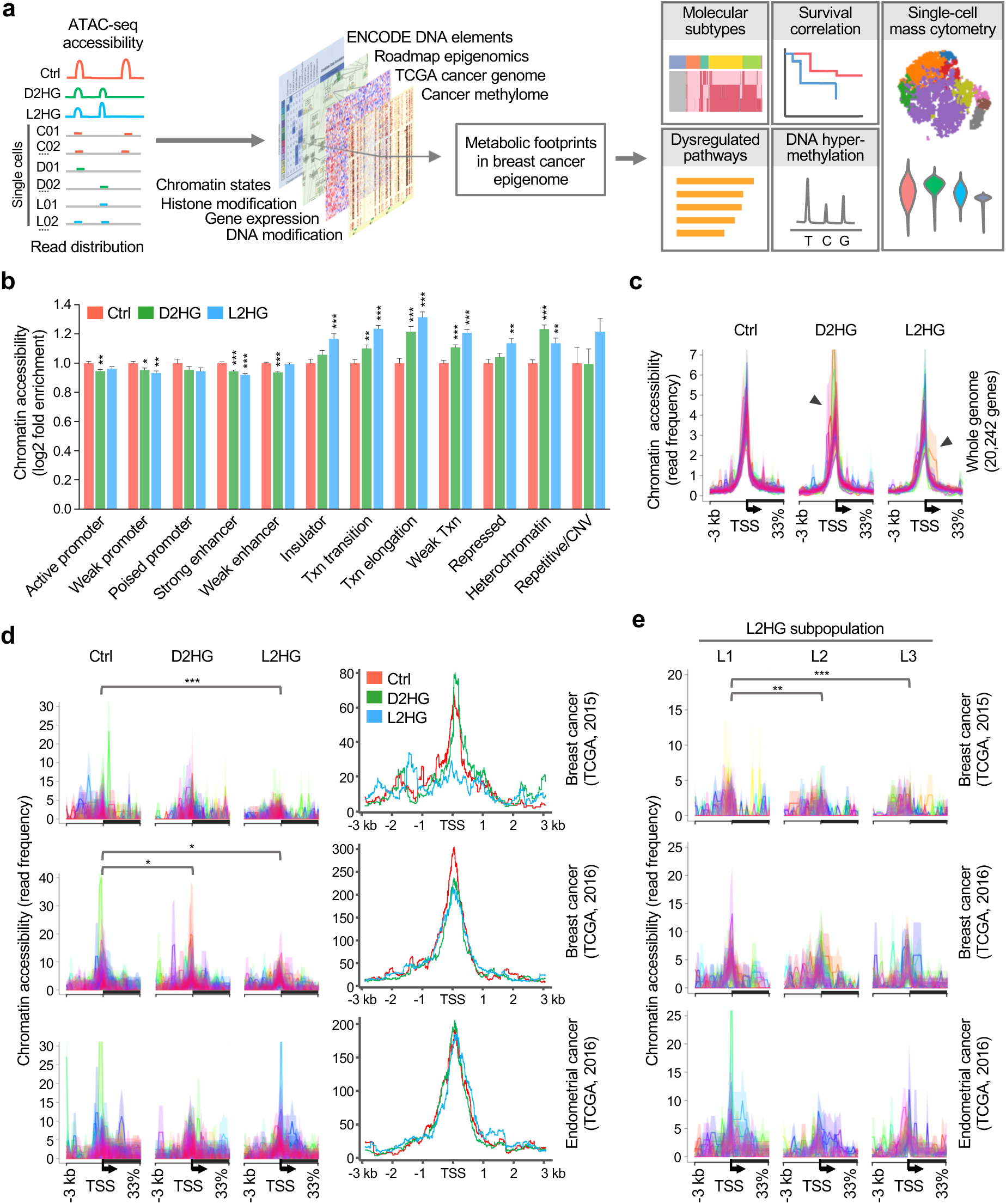
2HG decreases chromatin accessibility at genes that are highly methylated in breast cancer patients. **a**, Workflow of the integration of scATAC-seq data with high-throughput omics datasets to characterize clinically relevant epigenetic signatures imposed upon 2HG exposure. **b**, Distribution of ATAC-seq signals across ENCODE-ChromHMM functionally annotated regions in response to 2HG perturbation. **c**, Metagene analysis of chromatin accessibility assayed by scATAC-seq. Epigenomic accessibility landscape is centered on TSS across the genome (n = 20,242 genes). Colors represent individual cells. Solid lines indicate mean values and semi-transparent shade around the mean curve shows s.e.m. across the region. While a sharp and abrupt peak is evident over TSS in unexposed control cells, accessibility peaks are distorted in 2HG-exposed cells (arrowheads). **d**, Chromatin accessibility profiles for highly methylated genes in the TCGA cancer cohorts. scATAC-seq profiles in 2HG-exposed and unexposed cells are centered on TSS of highly methylated genes (n = 150 genes) in breast cancer (two independent datasets) and endometrial cancer patients. Colors represent single cells (left) and average accessibility profiles in each experimental condition (right). **e**, Chromatin accessibility profiles in L2HG subpopulations across TSS of highly methylated genes (n = 150 genes) in the TCGA cancer cohorts. \**P* < 0.05, \*\**P* < 0.01, \*\*\**P* < 0.001 versus unexposed control cells by one-way ANOVA followed by Dunnett’s multiple comparison test. ENCODE, Encyclopedia of DNA Elements; ChromHMM, chromatin Hidden Markov Modeling; TSS, transcriptional start site; TCGA, The Cancer Genome Atlas; Txn transition, transcriptional transition; Txn elongation, transcriptional elongation; Weak Txn, weak transcribed; CNV, copy number variation.

To address the sparse nature of scATAC-seq data^37,38^, we investigated regulatory variation in single-cell measurements by aggregating scATAC-seq signals over transcription start sites (TSS) in the genome. As shown in Fig. 3c with each color representing DNA accessibility around TSS regions (n = 20,242) in a single cell, the chromatin occupancy pattern in untreated control cells was highly concordant whereas the peaks were found to be diffused and heterogeneous in exposed cells, suggesting enhanced epigenetic variability in regulatory modules imposed by 2HG. However, the differences between exposed and unexposed cells were not quantitatively remarkable when chromatin accessibility was assessed over all TSS regions across the entire genome. In contrast, a marked reduction in accessibility was detected when a set of highly methylated genes (n = 150) in breast tumors^44^ was investigated (Fig. 3d and Supplementary Fig. 3b). Similar results were obtained using genes that were independently found to be hypermethylated in promoter regions^28^ (n = 150), indicating that 2HG led to reduced chromatin accessibilities in gene promoters displaying DNA hypermethylation in breast cancer. In contrast, no significant accessibility changes were observed in sets of genes that were highly methylated in HMECs (data not shown) or other tumor types including endometrial cancer. Next, we investigated epigenetic variability among the three L2HG subpopulations and found that L2 and L3 subsets exhibited a significant decrease in accessibility in comparison to L1 subset, which is consistent with earlier observations that L1 may represent a cell subpopulation that is potentially less responsive to oncometabolite perturbation (Fig. 3e). Collectively, single-cell chromatin profiling suggests that 2HG disrupts the fine-tuned spatial control of the mammary epigenome and reshapes the chromatin accessibility atlas of regulatory modules, potentially leading to DNA methylator phenotype.

### 2HG-mediated chromatin remodeling is linked to transcriptional repression and adverse prognosis associated with tumor hypermethylation

To assess the clinical relevance of tumor-associated hypermethylated genes that exhibited concomitant chromatin compaction in response to 2HG, we first analyzed RNA-seq whole-transcriptome profiles of 521 breast tumors and 112 adjacent uninvolved tissues from the TCGA cohort (Fig. 4a). Upon unsupervised hierarchical clustering, the majority of highly methylated genes appeared to be downregulated in all five PAM50 molecular subtypes, although no statistical significance was observed in luminal A tumors (Fig. 4a and Supplementary Fig. 4a). Strikingly, transcriptional repression was most evident in basal-like breast cancer, 84% of which displayed a triple-negative phenotype (i.e., negative for expression of estrogen, progesterone and HER2/neu receptors). These findings are in agreement with earlier results showing that 2HG levels were elevated predominantly in ERα-negative or basal-like breast cancer. In addition, none of the breast cancer patients investigated had 2HG-producing mutations in either the *IDH1* or *IDH2* gene (Fig. 4a).

**Fig. 4.**
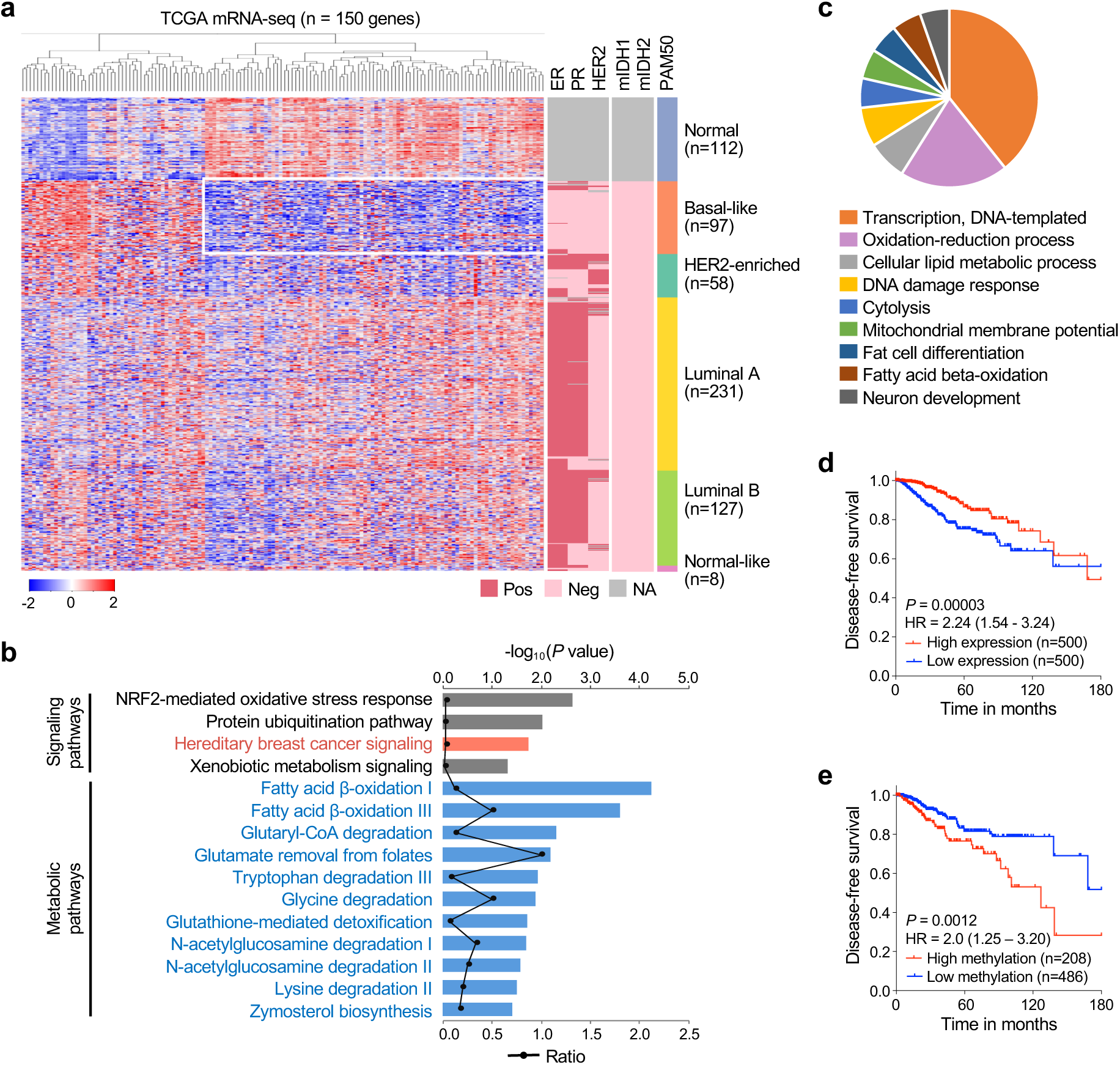
Tumor-associated DNA hypermethylation is linked to transcriptional repression and poor patient prognosis in basal-like breast cancer. **a**, Heat map depicting mRNA expression of highly methylated genes whose promoter regions showed diminished chromatin accessibility following 2HG exposure. The column dendrogram indicates unsupervised hierarchical clustering of 150 genes that are highly methylated in the TCGA breast cancer cohort. Rectangle outlined in white represent genes downregulated in basal-like tumors. Vertical sidebars indicate the category of each patient with regard to *IDH1/2* mutations (mIDH1/2), ER, PR, HER2 and PAM50 status. **b**, Overrepresented canonical signaling and metabolic pathways from IPA profiling of highly methylated genes in the TCGA breast cancer cohort. Colored bars (top x axis) represent −log_10_ *P* values obtained by Fisher’s exact test whereas the black line (bottom x axis) indicates the ratio between the number of genes compared to the total number of genes in a given pathway. **c**, Pie chart indicating GO biological processes enriched in genes associated with tumor hypermethylation. See also Supplementary Table 2. **d**,**e**, Prognostic significance of concurrent transcriptional repression (**d**) and hypermethylation (**e**) of genes shown in Supplementary Figure 4d. Disease-free survival (DFS) was investigated using Kaplan-Meier analysis, and log-rank (Mantel-Cox) *P* values and hazard ratios (HR) are shown (see Methods for further details). TCGA, The Cancer Genome Atlas; ER, estrogen receptor; PR, progesterone receptor; HER2, human epidermal growth factor receptor 2; PAM50, prediction analysis of microarray 50; IPA, Ingenuity Pathways Analysis; GO, Gene Ontology.

We next adopted functional pathway enrichment analyses to examine the physiological role of tumor-associated DNA hypermethylation. Ingenuity pathway analysis (IPA)^45^ revealed enrichment of genes involved in hereditary breast cancer signaling, reproductive system disease and gene expression (Fig. 4b and Supplementary Fig. 4b). In line with IPA analysis, 39% of the highly methylated genes were linked to DNA-templated transcription in Gene Ontology (GO) biological processes (Fig. 4c). In addition to DNA damage response (DDR), the most enriched GO pathways included lipid metabolic process, oxidation-reduction, fat cell differentiation and fatty acid beta-oxidation, all of which were relevant to intracellular metabolic signaling (Supplementary Fig. 4c and Supplementary Table 2), and therefore suggest an intimate entwinning of metabolic derangements with DNA hypermethylation in breast cancer.

To investigate whether altered expression of hypermethylated genes can impact the clinical outcome in breast cancer patients, we performed survival analysis of the TCGA breast cancer cohort. Approximately 10% of the genes exhibited a significant association between low expression and shorter disease-free survival (DFS) (Supplementary Fig. 4d). Strikingly, their combination displayed a steeper drop in survival (HR = 2.24; 95% CI, 1.54 to 3.24; *P* = 0.00003), compared to when the genes were analyzed as single variables (Fig. 4d). This suggests that concurrent downregulation of hypermethylated genes may have an additive adverse effect on patient prognosis. We further assessed the outcome of tumor hypermethylation on patient survival and found that the patients with a concurrent hypermethylation signature suffered decreased DFS (Fig. 4e). Together, the results suggest that 2HG-mediated loss of promoter accessibility, DNA methylator phenotype and concurrent transcriptional repression may present a high competing risk of mortality in breast cancer patients.

### 2HG enantiomers initiate tumor-associated promoter hypermethylation

To investigate whether transcriptional repression of hypermethylated genes is correlated with chromatin accessibility landscape, we leveraged the recently reported ATAC-seq data of 70 primary breast cancers from the TCGA cohort^46^. We found that the hypermethylated genes with transcriptional repression (boxed in Fig. 4a) showed a decrease in chromatin accessibility in basal-like or HER2-enriched breast cancers in comparison to luminal A, luminal B and normal-like tumors (Fig. 5a,b). We next sought to investigate if tumor-associated chromatin landscape could be attributable to 2HG-mediated epigenetic remodeling. To this end, we analyzed open chromatin signals corresponding to ChromHMM-annotated active promoter regions. The scATAC-seq peaks in unexposed control cells were comparable with DNase-seq and H2AFZ/H2A.Z signals that are localized at gene promoters facilitating RNA polymerase II occupancy^47^, while accessibility peaks were significantly reduced in D2HG- or L2HG-exposed cells (Fig. 5c). This finding suggests that oncometabolites could induce a decrease in chromatin accessibility at promoter regions of highly methylated genes that are downregulated in breast cancer. Additionally, whole-genome bisulfite sequencing (WGBS) data indicated that accessible promoter regions were essentially devoid of DNA methylation in control HMECs. These observations prompted us to investigate whether 2HG can initiate tumor-associated promoter hypermethylation accompanied by restricted chromatin accessibility evident in breast tumors.

**Fig. 5.**
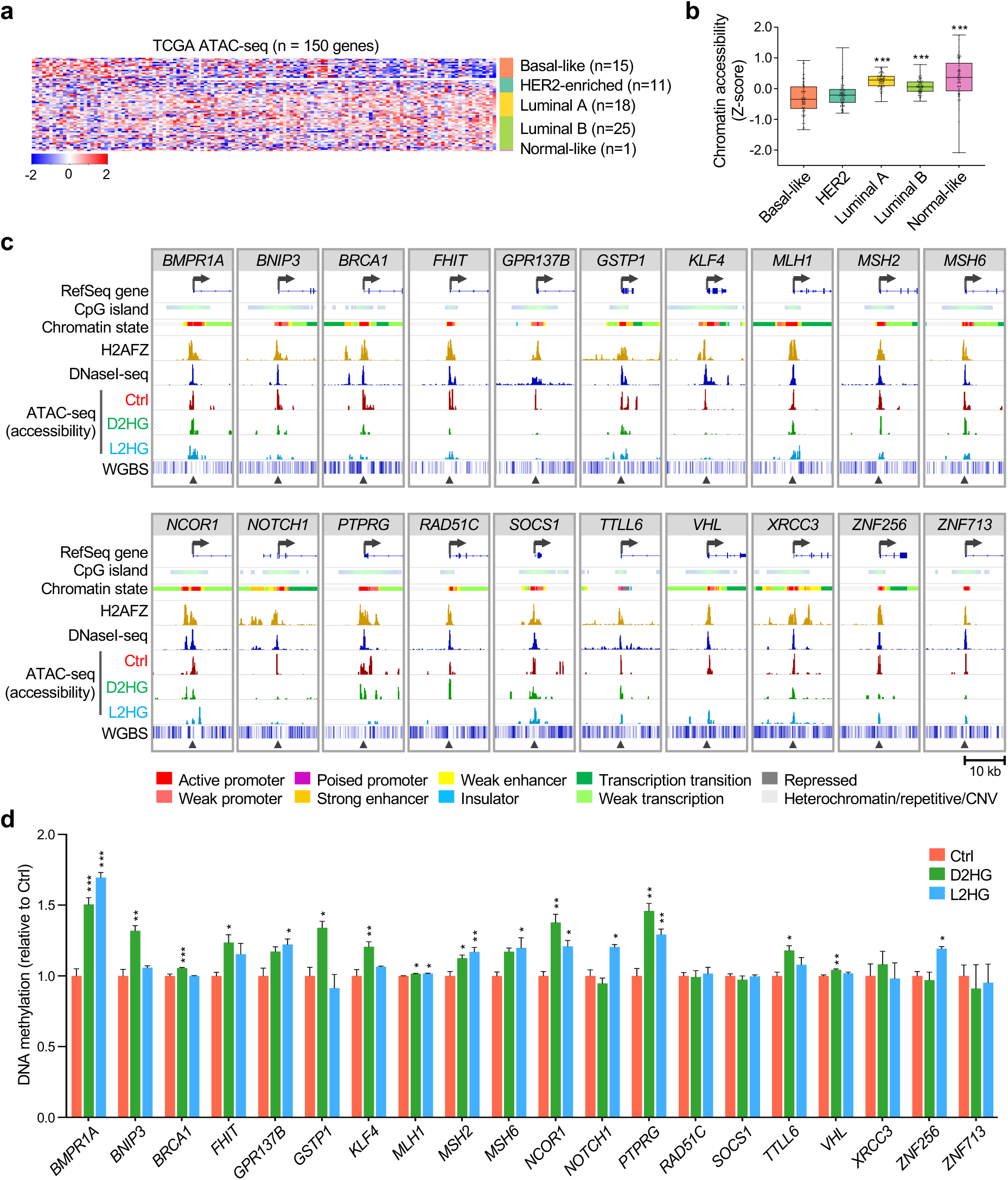
2HG induces DNA hypermethylation of tumor suppressor genes accompanied by loss of promoter accessibility. **a**, Heat map showing a decrease in chromatin accessibility of hypermethylated genes (n = 150) in basal-like and HER2-enriched breast cancer subtypes. Rectangle outlined in white represents genes downregulated in basal-like tumors as indicated in Fig. 4a. Vertical sidebar indicates five different molecular subtypes. **b**, Quantitative comparison of chromatin accessibility of hypermethylated genes across the TCGA breast cancer subtypes. The upper and lower whiskers indicate the minimum and maximum values of the data, center lines indicate the median, and the first and third quartiles are indicated by the bottom and top edges of the boxes respectively. \*\*\**P* < 0.001 versus basal-like tumors by one-way ANOVA with Dunnett’s multiple comparison test: basal-like (n = 15), HER2-enriched (n = 11), luminal A (n = 18), luminal B (n = 25) and normal-like (n = 1) subtypes. **c**, IGV genome browser tracks showing diminished chromatin accessibility at gene promoters following 2HG exposure. ChromHMM-defined chromatin states (GSE38163), DNaseI-seq (GSE29692), H2AFZ/H2A.Z ChIP-seq (GSE29611), WGBS profiles (GSE86732) from HMECs are shown. Arrows indicate TSS and direction of transcription initiation, and y-axes show read coverage. Histone variant H2A.Z is enriched around TSS. **d**, Promoter methylation measured by oxBS pyrosequencing. \**P* < 0.05, \*\**P* < 0.01, \*\*\**P* < 0.001 versus control by one-way ANOVA with Dunnett’s multiple comparison test (n ≥ 3 individual replicates). Error bars denote s.e.m. Validation by MeDIP-qPCR was also done in the same samples and promoter hypermethylation following 2HG exposure was confirmed using hTERT-HME1 cells (see Supplementary Figure 5a,b). IGV, Integrative Genomics Viewer; ChromHMM, chromatin Hidden Markov Modeling; WGBS, whole-genome bisulfite sequencing; TSS, transcriptional start site; oxBS, oxidative bisulfite; MeDIP, methylated DNA immunoprecipitation.

To evaluate DNA methylation at gene promoters, we utilized oxidative bisulfite (oxBS) conversion followed by pyrosequencing. DNA methylation levels across promoter CpG sites of tumor suppressor genes including but not limited to DNA damage response (DDR) genes were elevated in 2HG-exposed cells (Fig. 5d). Specifically, a marked methylation gain was observed in the majority of genes examined including *BRCA1*, *MSH2* and *MLH1*. Promoter hypermethylation in response to 2HG was confirmed by methylated DNA immunoprecipitation followed by qPCR (MeDIP-qPCR) (Supplementary Fig. 5a). A similar, albeit less pronounced, accumulation of *de novo* DNA methylation was also seen in immortalized, non-transformed mammary epithelial cells (hTERT-HME1) (Supplementary Fig. 5b). To further address the disease relevance of 2HG-mediated promoter hypermethylation, we evaluated our breast cancer cohort consisted of 77 primary tumors and 10 uninvolved tissue specimens^48^. Methyl-CpG binding domain proteins followed by sequencing (MBDCap-seq) indicated accumulation of DNA methylation at target promoter regions in tumor samples in comparison with their normal counterparts (Supplementary Fig. 5c). Collectively, these data suggest that 2HG induces loss of promoter accessibility accompanied by DNA hypermethylation of tumor suppressor genes, which is a prominent hallmark of the breast cancer epigenome.

### Single-cell mass cytometry reveals epigenetic plasticity and phenotypic heterogeneity

To disentangle epigenetically heterogeneous responses to 2HG at single-cell resolution, we next performed high-dimensional mass cytometry^49,50^ on viably cryopreserved cell suspensions from 2HG-exposed or unexposed control HMECs as well as cells exposed to 2HG followed by 5-day withdrawal (Supplementary Fig. 6a,b). The panel of metal-conjugated antibodies was designed to detect both chromatin modifications and intracellular proteins including BRCA1, MSH2 and MLH1 that are involved in DDR and checkpoint signaling (Supplementary Fig. 6a). Consistent with mass spectrometry measurements (Fig. 1i and Supplementary Fig. 1e), mass cytometry analysis showed that 2HG exposure led to a substantial increase in all histone markers investigated (Supplementary Fig. 6c). The biaxial gating of live single cells indicated a concurrent increase in multiple distinct classes of histone methylation and the presence of cell populations that were potentially less responsive to 2HG oncometabolites (Supplementary Fig. 6c,d).

To further characterize cell-to-cell variability in the chromatin landscape, we next applied nonlinear dimensionality reduction using *t*-distributed stochastic neighbor embedding (*t*-SNE) analysis. The *t*-SNE visualization revealed that control and withdrawal groups had similar multidimensional phenotypes and cell density distributions, which were readily distinct from those of 2HG-perturbed cells (Fig. 6a). Accordingly, altered chromatin modifications were effectively reverted by 2HG withdrawal (Supplementary Fig. 6c,d), suggesting that chromatin remodeling imposed by 2HG is essentially reversible. Self-organizing maps generated by FlowSOM further identified six discrete clusters as shown in Fig. 6b-d. Strikingly, a large population of HMECs (Cluster 1) displayed a concomitant increase in histone markers following 2HG exposure and reciprocal decrease upon withdrawal (Fig. 6e).

**Fig 6.**
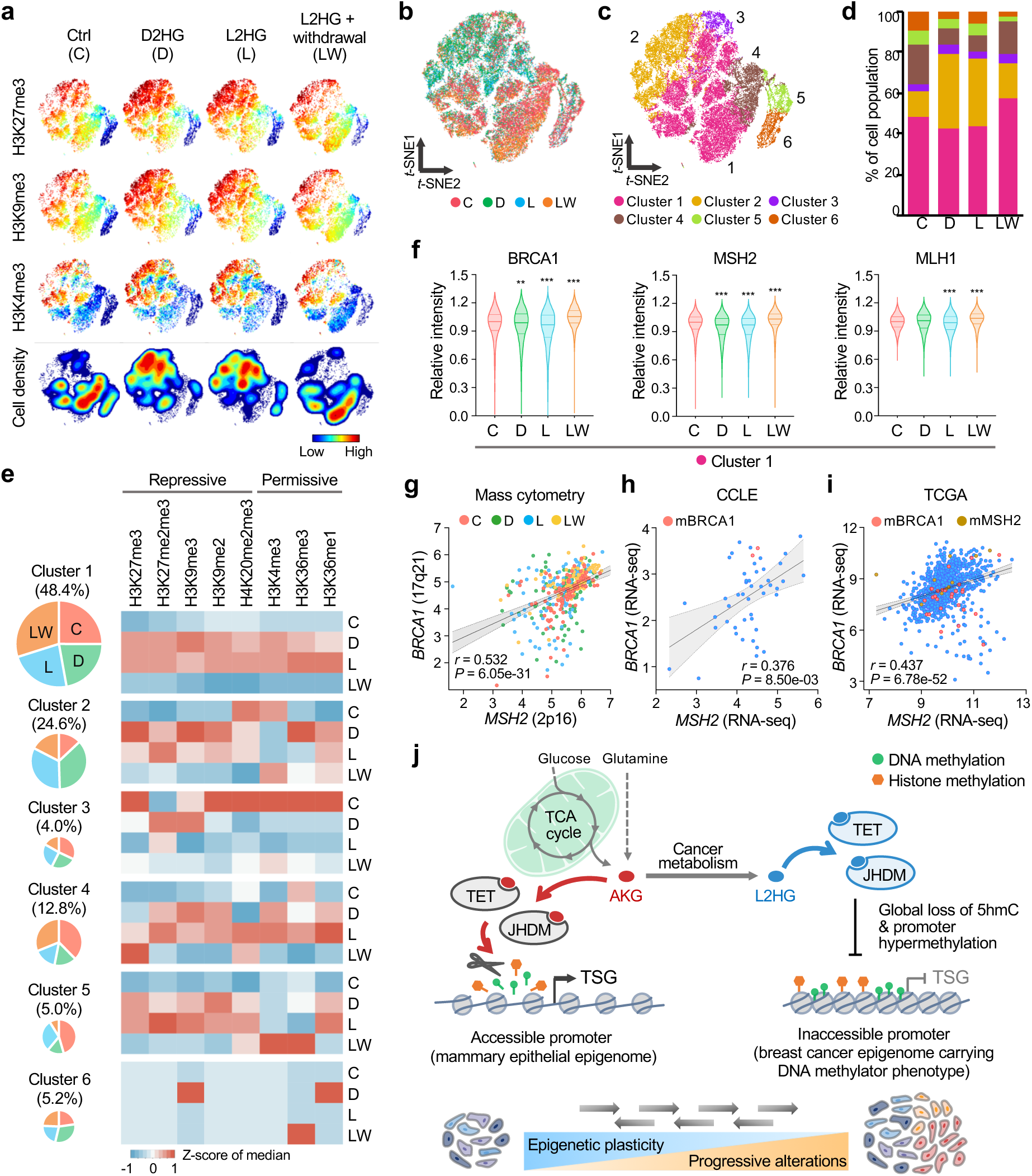
Single-cell mass cytometry demonstrates downregulation of hypermethylated genes in 2HG-responsive cell subpopulations. **a**, Representative *t*-SNE plots showing levels of histone modification markers in response to 2HG exposure. C, unexposed control cells; D, cells exposed to D2HG for 72 hr; L, cells exposed to L2HG for 72 hr; LW, cells exposed to L2HG for 72 hr followed by 5-day withdrawal of L2HG. Each data point on the *t*-SNE maps represents an individual cell and its color corresponds to cellular levels of each marker assessed. Density plots at the bottom show cell population distribution in each experimental group (~10,000 cells per condition). **b**, *t*-SNE plot of HMECs from all experimental groups merged. Each data point is colored by condition. **c**, *t*-SNE projections of epigenetically distinct cell subsets defined by FlowSOM. Cells are colored according to the cluster they were assigned to using consensus hierarchical clustering in FlowSOM analysis. **d**, Bar chart showing changes in cell frequency distribution across different treatment groups. **e**, Heat maps depicting epigenetically heterogeneous responses upon 2HG exposure. Normalized median values of signal intensities are shown for each cluster. Pie charts indicate the proportion of cells from different experimental groups in each cluster. **f**, Violin plots showing expression levels of the *BRCA1*, *MSH2* and *MLH1* genes in Cluster 1. \*\**P* < 0.01, \*\*\**P* < 0.001 versus control by one-way ANOVA followed by Dunnett’s multiple comparison test. **g**,**h**,**i**, Scatter plots showing the correlation between the expression of BRCA1 and MSH2 in the mass cytometry Cluster 1 (**g**), CCLE breast cancer cell lines (**h**) and TCGA breast cancer cohort (**i**). Data in **g** are transformed to arcsinh scales with the cofactor of 5 and four experimental groups (C, D, L and LW) are indicated by different colors. In the CCLE and TCGA datasets, red and brown dots indicate cancer cell lines or tumors with mutations in the *BRCA1* and *MSH2* genes respectively. Spearman’s correlation coefficient (*r* with *P* value) is indicated at the bottom of the panel. **j**, A model for metabolic rewiring of the breast cancer epigenome. TCA cycle metabolite αKG enhances TET and JHDM enzyme activities facilitating active DNA and histone demethylation such that promoter regions of tumor suppressor genes (TSG) remain accessible. Cancer metabolism accompanied with intratumor accumulation of 2HG oncometabolites, which antagonize DNA hydroxylases or histone demethylases, may drive global loss of 5hmC and promoter hypermethylation leading to DNA methylator phenotype. L2HG exposure could also confer enhanced cell-to-cell variability in the chromatin regulatory landscape by reversibly remodeling the breast cancer epigenome, potentially contributing to extensive intratumor heterogeneity. *t*-SNE, *t*-distributed stochastic neighbor embedding; FlowSOM, flow cytometry data analysis using self-organizing maps.

We next asked if 2HG-induced chromatin remodeling correlated with altered expression of DDR genes displaying promoter hypermethylation in response to 2HG (Fig. 5d). As expected, 2HG-responsive Cluster 1 cell subsets exhibited diminished expression of BRCA1, MSH2 and MLH1, which was restored upon withdrawal (Fig. 6f). In addition, BRCA1 downregulation was inversely correlated with H4K20 methylation (Supplementary Fig. 6e) and, to our surprise, we observed a remarkable association between the expression of the *BRCA1* and *MSH2* genes in Cluster 1 (Fig. 6g). Interestingly, this positive association was also evident in the CCLE cancer cell lines as well as in the TCGA breast cancer cohort (Fig. 6h,i). Although the precise mechanism is currently unknown, the findings may suggest that the two DDR genes are controlled by a shared set of transcriptional regulators, possibly via epigenetic modulators. Taken together, the multidimensional investigation of chromatin regulators directly implicates that the epigenome reprogramming imposed by 2HG and subsequent downregulation of DDR genes associated with tumor hypermethylation could contribute to the pathogenesis of breast malignancies with high 2HG accumulation.

## Discussion

Breast cancer cells accumulate high levels of 2HG with preferential concentration of L2HG in basal-like subtypes. Here we show that either of the 2HG enantiomers is sufficient to broadly confer two hallmarks of the breast cancer epigenome, i.e., global loss of 5hmC and promoter hypermethylation. Following 2HG exposure, promoter regions of the tumor suppressor genes involved in DDR signaling displayed reduced chromatin accessibility accompanied by methylation gain. The associated downregulation of BRCA1, MSH2 and MLH1 in 2HG-responsive cell subpopulations was validated by mass cytometry. Dysfunctional DDR pathway is one of the inherent characteristics of ‘BRCAness’, which is commonly seen in basal-like breast tumors and is manifested by an enhanced mutation rate and genomic instability^51^. Besides 2HG-mediated inhibition of active DNA demethylation, recent studies show that 2HG can directly inhibit αKG-dependent DDR signaling^19,52,53^ or indirectly alter expression of DNA repair genes^54,55^, suggesting that intratumor accumulation of 2HG potentially induces DDR deficiency via multiple signaling pathways. It is of interest to note in this context that 2HG exposure has been reported to establish the BRCAness phenotype in clinically pertinent models including patient-derived glioma cell lines and primary AML bone marrow cultures^55^. These findings may be relevant for designing treatment strategies for breast cancer patients with high intratumor 2HG, since effector pathways such as defective DDR signaling could represent an alternate targetable vulnerability in specific tumor subtypes with a stem cell-like transcriptional signature^30^, other than counteracting 2HG overproduction per se, for instance by using IDH small molecule inhibitors.

Aside from promoter hypermethylation, we show that 2HG inhibits a multitude of histone demethylases resulting in accumulation of both repressive and permissive chromatin marks. Further, genome-scale profiling of chromatin accessibility revealed selective loss of accessibility in active or poised chromatin and gain of accessibility in repressive or quiescent chromatin states in response to 2HG. These results indicate previously uncharacterized, two opposing effects of the oncometabolites imposed on the cellular epigenome, suggesting multifaceted entwinning of cancer metabolism with epigenetic regulation. Moreover, alterations in open chromatin occupancy were detected at enhancer and insulator regions. Of note, intratumor D2HG has been shown to dysregulate insulator or chromatin boundary function and promote aberrant gene-enhancer interaction, leading to constitutive activation of the glioma oncogene *PDGFRA*^56^. The findings therefore suggest that metabolic perturbations in breast cancer may modulate key aspects of chromatin functions including the three-dimensional (3D) genome topology.

Tumor heterogeneity is implicated in a wide range of neoplastic events involved in tumor evolution, dissemination, relapse or drug resistance^2^. Characterization of the molecular origin of intratumor diversity is thus integral to understanding and harnessing tumor heterogeneity and may provide new opportunities to define tumor subtypes or to design effective treatment. While genetic alterations have provided early insights into tumor heterogeneity, less attention has been paid to epigenetic diversity in breast cancer. In this study, scATAC-seq was applied to address cell-to-cell variability in the chromatin regulatory landscape and we found that the two enantiomers induced distinct chromatin accessibilities in the mammary epigenome. It is noted in this context that L2HG-exposed cells exhibited greater variability in comparison to D2HG-exposed cells, which is supported by the observations that hypoxic induction of L2HG could contribute to the development of epigenetic heterogeneity in glioblastoma^17^. Furthermore, single-cell profiling of epigenetic modifications by mass cytometry identified multiple cell subpopulations differentially responding to 2HG enantiomers. These observations suggest that metabolic dysfunction may enhance epigenetic cell-to-cell variability in chromatin regulatory modules, which can functionally and dynamically contribute to intratumor heterogeneity.

Of note, our findings and previous studies have suggested that high 2HG breast tumors without *IDH* mutations are associated with poor survival^30^, whereas a rare breast cancer subtype carrying *IDH* mutations has been reported to show better prognosis^34,35^. Consistently, prolonged survival has been observed in glioblastoma and anaplastic astrocytoma patients with *IDH* mutations^57,58^. These observations imply that the pathological mechanisms of high 2HG tumors may vary in the presence or absence of *IDH* mutations, which can be partly ascribed to the different potencies of the two enantiomers, i.e., L2HG is relatively more potent than D2HG that is produced by mutated IDH enzymes^18–20^. Alternatively, L2HG induction has been shown to be more sensitive to metabolic disturbances and generated via promiscuous enzymatic activity under hypoxic^16,17^ or acidic conditions^14,15^, which may yield cumulative adverse impacts on patient prognosis. Additionally, L2HG has been shown to promote the development of epigenetic cell-to-cell variability^17^. Intriguingly, more recent studies indicate that JHDM histone demethylases can directly sense cellular oxygen levels to regulate cell fate decisions^59,60^. In analogy, it might be plausible that αKG-dependent dioxygenases including TET DNA hydroxylases and JHDM histone demethylases could serve as metabolic sensors by detecting intratumor 2HG levels, which may reflect profound metabolic disturbances accompanied by underlying acidosis and hypoxia implicated in the tumor microenvironment.

Collectively, the present study substantiates that 2HG imposes metabolic footprints on the mammary epithelial epigenome by impairing DNA and histone demethylation, and provides a molecular basis in which altered metabolism leads to loss of 5hmC and associated promoter hypermethylation, thereby priming early epigenetic events to be exploited during breast cancer development (Fig. 6j). This in turn suggests that defective metabolic fluxes can disrupt epigenetic homeostasis resulting in loss of chromatin integrity. Nonetheless, 2HG-induced epigenetic liabilities are found to be dynamic and essentially reverted upon withdrawal, highlighting the inherent plasticity of the cellular epigenome. Finally, our findings suggest that chromatin and its regulatory modules are highly vulnerable to intracellular metabolic cues and future investigation on metabolism-epigenome coupling may lead to the identification of key molecular signatures that determine breast cancer susceptibility.

## Methods

### Cell lines, growth conditions and reagents

Human breast cancer cell lines BT20, BT474, MCF7, MDA-MB-157, MDA-MB-231 and MDA-MB-361, and non-malignant immortalized epithelial cells hTERT-HME1, 184B5 and MCF12A were acquired from the American Type Culture Collection (ATCC). Unless otherwise stated, cancer cell lines were maintained in DMEM (Gibco) supplemented with 10% fetal bovine serum (Sigma-Aldrich) and 100 U/ml penicillin plus 100 μg/ml streptomycin (Gibco) as previously reported^61,62^. Primary HMECs were obtained from Invitrogen and maintained at low passage number (below 5). HMECs, hTERT-HME1, 184B5 and MCF12A cells were cultured in mammary epithelial growth medium according to the manufacturer’s instructions. Authentication of cell line genomic DNA was performed at ATCC using DNA fingerprint analysis of polymorphic, short tandem repeat sequences. Exposure to cell-permeable 2HG analogues was carried out by supplementing octyl esters of *R*-2-hydroxyglutarate or *S*-2-hydroxyglutarate (Cayman Chemical) to the culture medium at a final concentration 72 hr before harvesting. Dimethyloxalylglycine (DMOG) was obtained from Sigma-Aldrich. Culture medium was replaced daily with fresh complete medium with or without oncometabolite supplementation.

### Metabolite extraction and quantification by LC-MS

Following dissociation, cells were washed twice with ice-cold phosphate-buffered saline (PBS) and cell pellets were flash-frozen on dry ice. For αKG analysis, metabolites were extracted with 80:20 methanol:water (−80°C) containing stable isotope-labeled internal standard [1,2,3,4-^13^C_4_]α-ketoglutaric acid (Cambridge Isotope Laboratories) and incubated at −80°C for 1 hr as described previously^63,64^. Extracts were then centrifuged at 13,800*g* for 10 min and supernatants were transferred to glass autosampler vials for high-performance liquid chromatography-electrospray ionization-mass spectrometry (HPLC-ESI-MS) measurements. For 2HG analysis, cells were processed as mentioned above except that [1,2,3,4-^13^C_4_]L-malic acid (Cambridge Isotope Laboratories) was added as an internal standard and dried extracts were derivatized with diacetyl-L-tartaric anhydride (DATAN, Sigma-Aldrich). HPLC-ESI-MS detection was conducted on a ThermoFisher Q Exactive mass spectrometer with on-line separation by a ThermoFisher Dionex Ultimate 3000 HPLC. HPLC conditions for αKG analysis were: column, Synergi Polar-RP, 4 μm, 2×150 mm (Phenomenex); mobile phase A, 0.1% formic acid in water; mobile phase B, 0.1% formic acid in acetonitrile; flow rate, 250 μl/min; gradient, 1% B to 5% B over 5 minutes, 5% B to 95% B over 1 min and held at 95% B for 2 min. HPLC conditions for 2HG analysis were: column, Luna NH2, 3 μm, 2×150 mm (Phenomenex); mobile phase A, 5% acetonitrile in water containing 20 mM ammonium acetate and 20 mM ammonium hydroxide, pH 9.45; mobile phase B, acetonitrile; flow rate, 300 μl/min; gradient, 85% B to 1% B over 10 min and held at 1% B for 10 min. The conditions used to selectively quantify D2HG and L2HG were: column, Kinetex C18, 2.6 μm, 2.1×100 mm (Phenomenex); mobile phase, 1% acetonitrile with 125 mg/l ammonium formate, pH 3.6; flow rate, 400 μl/min. Full scan mass spectra were acquired in the orbitrap using negative ion detection over a range of m/z 100–800 at 70,000 resolution (m/z 300). Metabolite identification was based on accurate mass match to the library ±5 ppm and agreement with the HPLC retention time of authentic standards. Quantification of metabolites was carried out by integration of extracted ion chromatograms with the corresponding standard curves.

### Immunofluorescence staining

Cells were plated in 8-well chamber slides (Falcon) at a density of 1-2×10^4^ cells/well at least 24 hr prior to 2HG exposure. Cell were then fixed with 4% paraformaldehyde (PFA) in PBS for 10 min at room temperature (RT) and permeabilized with 0.2% Triton X-100 in PBS for 10 min at RT. For 5hmC staining, permeabilized cells were treated with 2N HCl for 30 min at RT and neutralized with 100 mM Tris-HCl (pH 8.5). Nonspecific binding was blocked with 10% goat serum in 0.2% Triton X-100 and PBS for 1 hr at RT and stained with primary antibodies in PBS with 5% goat serum and 0.2% Triton X-100 overnight at 4°C. After incubation with Alexa Fluor-conjugated secondary antibodies (Molecular Probes) for 1 hr at RT, nuclei were stained for 5 min with DAPI (Sigma-Aldrich). Single optical sections were acquired using a Zeiss LSM710 confocal microscope and image quantification was performed with NIH ImageJ software (version 1.52n). Primary antibodies included rabbit polyclonal anti-5hmC (1:500; 39769, Active Motif) and rabbit monoclonal anti-H3K27me3 (1:800; 9733, Cell Signaling Technologies).

### Tet-assisted bisulfite (TAB) pyrosequencing

TAB pyrosequencing was used to differentiate 5hmC from 5mC^65^. High molecular weight genomic DNA was extracted using Gentra Puregene reagents (Qiagen), followed by an additional ethanol precipitation and resuspension in low-EDTA TE buffer (10 mM Tris-HCl, 0.1 mM EDTA, pH 8.0). RNase A and proteinase K digestion were included in the isolation procedure. UV absorbance was measured on a NanoDrop 2000 (ThermoFisher) and each DNA sample was routinely examined by agarose gel electrophoresis with GelRed staining to ensure the absence of contaminating RNA and degradation of genomic DNA. Isolated genomic DNA was then subjected to Tet-assisted bisulfite (TAB) treatment as we previously described^66^. After bisulfite conversion using EpiTect Fast Bisulfite Conversion kit (Qiagen), pyrosequencing was conducted on a PyroMark Q96 MD instrument using CpG LINE-1 assay (973043, Qiagen). To monitor bisulfite conversion efficiency, a C outside a CpG site was added within dispensation order for the sequence to be analyzed as a built-in control. The quantitative levels of 5mC and 5hmC for each CpG dinucleotide were determined using PyroMark CpG software (version 1.0, Qiagen).

### Multiplexed chromatin profiling by mass spectrometry

Nuclei were isolated from 2×10^6^ cells using Nuclear Isolation Buffer (NIB) composed of 15 mM Tris-HCl (pH 7.5), 60 mM KCl, 15 mM NaCl, 5 mM MgCl_2_, 1 mM CaCl_2_, 250 mM sucrose, 0.3% NP-40, 1 mM DTT plus 10 mM sodium butyrate added immediately prior to use, for 30 min on ice. Nuclei were pelleted at 600*g* for 5 min at 4°C and detergent was removed by washing twice with NIB without NP-40. Histones from isolated nuclei were acid extracted with 5 volumes of 0.2 M H_2_SO_4_ for 1 hr at RT. Cellular debris was removed by centrifugation at 4,000*g* for 5 min. Trichloroacetic acid was added to the supernatant at a final concentration of 20% (v/v) and incubated for 1 hr to precipitate histone proteins. Histones were pelleted at 10,000*g* for 5 min, washed once with 0.1% HCl in acetone, twice with 100% acetone followed by centrifugation at 15,000*g* for 5 min, and then briefly air-dried. Histones were derivatized, digested and analyzed by targeted LC-MS/MS as described previously^67–69^.

### Flow cytometry

Cell cycle phase distribution was analyzed by flow cytometry. Cells were fixed with ice-cold 70% methanol for 1 hr on ice. Following centrifugation, cells were washed with PBS and stained with 10 μg/ml propidium iodide (Sigma-Aldrich) solution in PBS containing 1 μg/ml DNase-free RNase A (ThermoFisher) and incubated in the dark on ice for 1 hr. Samples were then processed on a BD FACSCalibur flow cytometer equipped with CellQuest Pro software (version 5.2.1, Becton Dickinson) and data were analyzed using FlowJo software (version 7.6.5, Tree Star).

### Single-cell ATAC-seq library preparation

Single-cell ATAC-seq libraries were prepared on a Fluidigm C1 workstation using ‘ATAC Seq-Cell Load and Stain Rev C’ script as previously described^37^ with modifications. Briefly, cells were passed through a 20 μm cell strainer (CellTrics, Sysmex Partac) to remove debris and remaining cell aggregates and mixed at a ratio of 7:3 with C1 suspension reagent. The resulting single-cell suspension was loaded on C1 Single-Cell Open App IFC chip (1862x, 10-17 μm, Fluidigm) at a concentration of 350 cells*/*μl. Captured cells were stained with 2 μM green-fluorescent calcein-AM and 4 μM red-fluorescent ethidium homodimer-1 (Molecular Probes) and visualized under an EVOS FL cell imaging station (Life Technologies) to ensure successful capture and to determine cell viability. The single-cell capture rates were typically >80% and >90% of captured single cells were alive. After cell lysis and Tn5 transposition, 8 cycles of pre-amplification were run on IFC chip. Pre-amplified PCR products were transferred to 96-well plates and further amplified for an additional 13 cycles using custom Nextera dual-index primers and NEBNext High-Fidelity 2X PCR master mix (New England Biolabs). Individually barcoded libraries were pooled and purified on a single MinElute column (Qiagen). The quality and size distribution of pooled libraries were evaluated on an Agilent 2100 Bioanalyzer using High Sensitivity DNA reagents (Agilent).

### Single-cell ATAC-seq data analysis

scATAC-seq libraries were sequenced on a NextSeq 500 platform with High Output reagents (Illumina) using paired-end 75-bp reads. All scATAC-seq data were preprocessed as essentially described^37^. In short, adapter and primer sequences were trimmed and initial quality control checks were performed using FastQC tools (https://www.bioinformatics.babraham.ac.uk/ projects/fastqc/). Sequencing reads were aligned to the GRCh37/hg19 assembly of the human genome using Bowtie2 (ref. ^70^) with the parameter ‘-X2000’ to ensure paired reads were within 2 kb of one another. PCR duplicates were eliminated using Picard tools (version 2.9.2, http://broadinstitute.github.io/picard/) and alignments with mapping quality less than 30 were subsequently removed by samtools. Reads mapped to the mitochondria and unmapped contigs were filtered out and excluded from further analysis. PCA projections of scATAC-seq profiles were performed using SCRAT^71^, and gene feature was applied to aggregate sequencing reads from each cell, in which 3,000 bp upstream to 1,000 bp downstream of TSS is regarded as the region of interest for each gene. After aggregation, the signals for each feature were normalized to adjust for library size and model-based clustering (mclust) module was utilized to identify cell subpopulations. Peak calling was performed using MACS2 with the following settings: --nomodel --nolambda --keep-dup all --call-summits. Artifact signals were excluded using ENCODE blacklist^72^. Circular visualization of ATAC-seq signals was carried out by employing Circos tools (version 0.69-6, http://circos.ca).

### Breast cancer cohorts, resources and data analysis

Level 3 TCGA Breast Invasive Carcinoma (BRCA) data of tumor and normal samples were accessed from the Broad GDAC Firehose (http://gdac.broadinstitute.org) and RSEM-normalized RNA-seq values were log2 transformed before analysis. Unsupervised hierarchical clustering was utilized to distinguish mRNA expression profiles among different genes and heat maps were generated using heatmap.2 function implemented in gplots package of R statistical program. Clinical data including PAM50 intrinsic subtypes, ER/PR/HER2 expression and IDH mutation status were retrieved using the Cancer Genomics cBioPortal^73^ and were integrated into RNA-seq heat map. TCGA DNA methylation data generated using Infinium Human Methylation 450K (HM450K) BeadChip array were retrieved from the cBioPortal database. Normalized methylation scores at each CpG dinucleotide are expressed as *β* values, representing a continuous measurement from 0 (completely unmethylated) to 1 (completely methylated). In the event of multiple CpG probes per gene, the most negatively correlated with mRNA expression was selected. Chromatin accessibility data of TCGA primary tumor tissue samples were extracted from the UCSC Xena browser (https://xenabrowser.net/). After z-scale normalization of ATAC-seq signals, open chromatin occupancies at promoter regions were correlated with PAM50 gene signature to evaluate DNA accessibility profiles across breast cancer subtypes. Our breast cancer methylome data generated using MBDCap-seq are available at The Cancer Methylome System (http://cbbiweb.uthscsa.edu/KMethylomes/). Global chromatin profiling and metabolomics datasets were retrieved from the Broad Institute CCLE (https://portals.broadinstitute.org/ccle).

### Oxidative bisulfite (oxBS) pyrosequencing

To selectively detect 5mC modification, genomic DNA was subjected to oxBS conversion^74^ using TrueMethyl oxBS module (NuGEN Technologies) as per the manufacturer’s recommendations. In short, genomic DNA was affinity-purified using 80% acetonitrile (Fisher Scientific) and TrueMethyl magnetic beads to eliminate potential contaminating compounds. After the denaturation step, genomic DNA was oxidized to convert 5-hydroxymethylcytosine to 5-formylcytosine. Bisulfite treatment was then carried out to convert 5-formylcytosine to uracil, leaving 5-methylcytosine intact. Following desulfonation and purification, converted DNA was quantified using Qubit ssDNA assay (Invitrogen). PCR amplification of oxBS converted DNA was carried out with biotin-labeled primers. Primer design was carried out using PyroMark Assay Design software (version 2.0, Qiagen). Pyrosequencing of biotinylated PCR products was performed using PyroMark Q48 Advanced CpG reagents (Qiagen) on a Pyromark Q48 Autoprep apparatus (Qiagen) following the manufacturer’s protocol. 5mC levels at CpG sites were determined using PyroMark Q48 Autoprep software (version 2.4.2, Qiagen) in CpG Assay mode. All samples were prepared, amplified and sequenced in triplicates. PCR and pyrosequencing primers are listed in Supplementary Tables 3 and 4.

### Methylated DNA precipitation PCR (MeDIP-qPCR)

Prior to the 5mC immune-capture procedure, genomic DNA was fragmented to an average length of 200-600 bp using a Covaris 220 system. MeDIP was performed using MeDIP reagents (Active Motif) as per the manufacturer’s instructions. In brief, fragmented DNA was heat-denatured and immunoprecipitated with anti-5mC antibody (39649, Active Motif). An additional quantity of fragmented DNA equivalent to 10% of DNA being used in the immunoprecipitation reaction was also denatured and saved as input DNA. Immunoprecipitated DNA and input DNA were then purified with phenol/chloroform extraction and amplified using GenoMatrix Whole Genome Amplification kit (Active Motif). Quantitative PCR was performed using PowerUP SYBR Green master mix on an ABI StepOnePlus real-time PCR instrument (Applied Biosystems). All PCR reactions were run in triplicates. The relative enrichment of target sequences after MeDIP was evaluated by calculating the ratios of the signals in immunoprecipitated DNA versus input DNA. Locus-specific primers were designed with NCBI Primer-BLAST and synthesized by Integrated DNA Technologies. Primer sequences are provided in Supplementary Table 5.

### Panel design and heavy-metal labeling of antibodies

Prior to antibody conjugation, the antibody panel was designed by allocating targets to specific heavy-metal isotopes depending on the sensitivity of the mass cytometer, e.g., assigning low abundance targets to high sensitivity channels in order to minimize potential spectral overlap^75^. Subsequently, in-house conjugation of antibodies was performed using Maxpar X8 antibody labeling reagents (Fluidigm) as previously described^76^ with some modifications. Briefly, up to 100 μg of carrier-free IgG antibody was subjected to buffer exchange by washing with R-buffer using a 50 kDa Amicon filter (Millipore) that was pre-soaked with R-buffer. Antibodies were then partially reduced with 4mM TCEP (ThermoFisher) for 30 min at RT followed by washing with C-buffer. In parallel, metal chelation was carried out by adding lanthanide metal solutions (Fluidigm) to chelating polymers (Fluidigm) in L-buffer. Metal-loaded polymers were then washed with L-buffer and concentrated on a 3 kDa Amicon filter (Millipore). Partially reduced antibodies were incubated with metal-loaded polymers for 90 min at RT followed by washing with W-buffer. Following conjugation, antibody concentration was determined by spectrometry with a NanoDrop 2000 (ThermoFisher). Metal-conjugated antibodies were stored in antibody stabilization solution (Candor Bioscience) supplemented with 0.05% sodium azide at 4°C. The panel of metal-conjugated antibodies is provided in Supplementary Table 6.

### Multidimensional chromatin profiling by mass cytometry

Cell suspensions were prepared at a concentration of 1×10^7^ cells/ml in serum-free, protein-free medium and stained with 1 μM cisplatin (195Pt) for 5 min at RT to determine cell viability. After quenching with CyTOF buffer composed of PBS with 1% BSA (Invitrogen), 2mM EDTA (Ambion) and 0.05% sodium azide (Teknova), staining with lanthanide-conjugated antibodies was performed as previously described^50^, but with the following modifications. In brief, following extracellular marker staining, cells were fixed with 1.6% PFA (Electron Microscopy Sciences) for 15 min at RT and permeabilized with ice-cold methanol (Fisher Scientific) for 30 min at 4°C. After adding F_c_ receptor blocker (BioLegend), cells were labeled overnight at 4°C with a cocktail of antibodies recognizing chromatin modifications or intracellular components. On the next day, excess of antibodies were washed off with CyTOF buffer and cells were stained with 250 nM 191/193Ir-containing DNA intercalator (Fluidigm) in PBS with 1.6% PFA for 30 min at RT. After resuspending in double-deionized water, samples were kept on ice. Immediately prior to acquisition, cells were prepared at a concentration of 0.2-1.0×10^6^ cells/ml in 0.1X EQ bead solution containing four element calibration beads (Fluidigm) and filtered through a 20 μm cell strainer (CellTrics, Sysmex Partac) to remove any potential aggregates. Cells were then acquired at a rate of 300-500 events/s using a Helios mass cytometer (Fluidigm) and CyTOF software (version 6.7) with noise reduction, a lower convolution threshold of 400, event length limits of 10-150 pushes, a sigma value of 3 and a flow rate of 0.030 ml/min.

### Mass cytometry data analysis

Data analysis was conducted using the cloud-based platform Cytobank^77^ and the statistical programming environment R. Following data acquisition, mass cytometry data were normalized using EQ calibration beads as previously described^78^. Bead-normalized data were then uploaded onto Cytobank platform to carry out initial gating and population identification using the indicated gating schemes (Supplementary Fig. 6b). For downstream analysis, live single cells were identified based on 140Ce bead, event length, DNA content (191Ir) and live/dead (195Pt) channels. Histograms and two-dimensional contour plots were generated to assess the global levels of chromatin modifications across the samples. Using an equal number of randomly selected live singlets from each sample, dimensionality reduction was implemented by *t*-SNE analysis with the following settings: perplexity = 60, theta = 0.5, iteration = 1,000. FlowSOM clustering was carried out on the same data using the standard parameters to quantify changes in cell subsets in an unbiased manner. The 2D coordinates of the *t*-SNE map were fed to FlowSOM analysis for population identification based on hierarchical consensus clustering.

Comparisons of chromatin modifications among the samples in each cluster were performed by generating heat maps in R using gplots package and median signal intensities extracted from Cytobank.

### Statistical analysis

Pairwise comparisons were carried out with a two-tailed unpaired Student’s *t*-test and multiple comparisons were assessed using a one-way ANOVA followed by Dunnett’s multiple comparison post-hoc test unless otherwise indicated in the figure legends. For Kaplan-Meier survival analysis, expression or methylation values were classified as high or low by using the median as a cutoff value and disease-free survival data was used to measure prognosis. Log-rank (Mantel-Cox) test was used to evaluate statistical differences and hazard ratio was reported with 95% confidence interval. Statistical analyses were performed using GraphPad Prism program (version 8.1). For all statistical analyses, differences of *P* < 0.05 were considered statistically significant. All quantitative data are presented as mean ± s.e.m. unless specified otherwise.

## Data availability

The raw and processed single-cell ATAC-seq data have been deposited in the National Center for Biotechnology Information (NCBI) Gene Expression Omnibus (GEO) and are available under accession GSE135412. The R code used in the study is available upon request from the authors. All other data described, analyzed and represented in the figures that support the findings of this study are available from the corresponding authors upon request.

## Acknowledgements

We thank all laboratory members for helpful discussions and technical assistance. We are grateful to the BioAnalytics and Single-Cell Core (BASiC) for single-cell analysis, Mass Spectrometry Core for metabolite mass spectrometry, Optical Imaging Facility for confocal imaging, Genome Sequencing Facility for next-generation sequencing and Flow Cytometry Shared Resource Facility at the University of Texas Health Science Center at San Antonio for flow cytometry, Northwestern Proteomics Core Facility for proteomics analyses. This study was supported by NIH grants U54CA217297, P30CA054174 and CPRIT grant RP150600. J.R. acknowledges funding from the US National Science Foundation (ABI 1565076). M.K. is a recipient of the CPRIT predoctoral fellowship (RP170345).

## Author contributions

T.H.M.H., K.M. and M.K. jointly conceived the project, designed the experiments, interpreted the results and prepared the manuscript with contributions from all co-authors. M.K. carried out the majority of experiments and data analysis. J.R., M.Z. and C.L.L assisted with scATAC-seq data analysis, computational modeling and statistical methods. K.M. and M.K. established the methods for chromatin mass cytometry. C.M.W, N.D.L and N.K assisted with validation of metal-tagged antibodies. All authors read and approved the manuscript.

## Correspondence and requests for materials

should be addressed to T.H.M.H or K.M.

## References

1. DeBerardinis, R. J., Lum, J. J., Hatzivassiliou, G. & Thompson, C. B. The biology of cancer: metabolic reprogramming fuels cell growth and proliferation. Cell Metab. 7, 11–20 (2008).

2. McGranahan, N. & Swanton, C. Clonal heterogeneity and tumor evolution: past, present, and the future. Cell 168, 613–628 (2017).

3. Warburg, O. On the origin of cancer cells. Science 123, 309–314 (1956).

4. Intlekofer, A. M. & Finley, L. W. S. Metabolic signatures of cancer cells and stem cells. Nat. Metab. 1, 177–188 (2019).

5. Linster, C. L., Van Schaftingen, E. & Hanson, A. D. Metabolite damage and its repair or pre-emption. Nat. Chem. Biol. 9, 72–80 (2013).

6. DeBerardinis, R. J. & Chandel, N. S. Fundamentals of cancer metabolism. Sci. Adv. 2, e1600200 (2016).

7. Ryan, D. G. et al. Coupling krebs cycle metabolites to signalling in immunity and cancer. Nat. Metab. 1, 16–33 (2019).

8. Losman, J. A. & Kaelin, W. G., Jr. What a difference a hydroxyl makes: mutant IDH, (R)-2-hydroxyglutarate, and cancer. Genes Dev. 27, 836–852 (2013).

9. Ye, D., Guan, K. L. & Xiong, Y. Metabolism, activity, and targeting of D- and L-2-hydroxyglutarates. Trends Cancer 4, 151–165 (2018).

10. Dang, L. et al. Cancer-associated IDH1 mutations produce 2-hydroxyglutarate. Nature 462, 739–744 (2009).

11. Ward, P. S. et al. The common feature of leukemia-associated IDH1 and IDH2 mutations is a neomorphic enzyme activity converting alpha-ketoglutarate to 2-hydroxyglutarate. Cancer Cell 17, 225–234 (2010).

12. Mishra, P. et al. ADHFE1 is a breast cancer oncogene and induces metabolic reprogramming. J. Clin. Invest. 128, 323–340 (2018).

13. Tang, X. et al. A joint analysis of metabolomics and genetics of breast cancer. Breast Cancer Res. 16, 415 (2014).

14. Nadtochiy, S. M. et al. Acidic pH Is a metabolic switch for 2-hydroxyglutarate generation and signaling. J. Biol. Chem. 291, 20188–20197 (2016).

15. Intlekofer, A. M. et al. L-2-Hydroxyglutarate production arises from noncanonical enzyme function at acidic pH. Nat. Chem. Biol. 13, 494–500 (2017).

16. Oldham, W. M., Clish, C. B., Yang, Y. & Loscalzo, J. Hypoxia-mediated increases in L-2-hydroxyglutarate coordinate the metabolic response to reductive stress. Cell Metab. 22, 291–303 (2015).

17. Intlekofer, A. M. et al. Hypoxia induces production of L-2-hydroxyglutarate. Cell Metab. 22, 304–311 (2015).

18. Xu, W. et al. Oncometabolite 2-hydroxyglutarate is a competitive inhibitor of alpha-ketoglutarate-dependent dioxygenases. Cancer Cell 19, 17–30 (2011).

19. Chowdhury, R. et al. The oncometabolite 2-hydroxyglutarate inhibits histone lysine demethylases. EMBO Rep. 12, 463–469 (2011).

20. Koivunen, P. et al. Transformation by the (R)-enantiomer of 2-hydroxyglutarate linked to EGLN activation. Nature 483, 484–488 (2012).

21. Pfeifer, G. P., Xiong, W., Hahn, M. A. & Jin, S. G. The role of 5-hydroxymethylcytosine in human cancer. Cell Tissue Res. 356, 631–641 (2014).

22. Jones, P. A., Issa, J. P. & Baylin, S. Targeting the cancer epigenome for therapy. Nat. Rev. Genet. 17, 630–641 (2016).

23. Laird, P. W. Cancer epigenetics. Hum. Mol. Genet. 14, R65–76 (2005).

24. Tsai, K. W. et al. Reduction of global 5-hydroxymethylcytosine is a poor prognostic factor in breast cancer patients, especially for an ER/PR-negative subtype. Breast Cancer Res. Treat. 153, 219–234 (2015).

25. Jin, S. G. et al. 5-Hydroxymethylcytosine is strongly depleted in human cancers but its levels do not correlate with IDH1 mutations. Cancer Res. 71, 7360–7365 (2011).

26. Yang, L., Yu, S. J., Hong, Q., Yang, Y. & Shao, Z. M. Reduced expression of TET1, TET2, TET3 and TDG mRNAs are associated with poor prognosis of patients with early breast cancer. PLoS One 10, e0133896 (2015).

27. Pfister, S. X. & Ashworth, A. Marked for death: targeting epigenetic changes in cancer. Nat. Rev. Drug Discov. 16, 241–263 (2017).

28. Thienpont, B. et al. Tumour hypoxia causes DNA hypermethylation by reducing TET activity. Nature 537, 63–68 (2016).

29. Skvortsova, K. et al. DNA hypermethylation encroachment at CpG island borders in cancer is predisposed by H3K4 monomethylation patterns. Cancer Cell 35, 297–314.e298 (2019).

30. Terunuma, A. et al. MYC-driven accumulation of 2-hydroxyglutarate is associated with breast cancer prognosis. J. Clin. Invest. 124, 398–412 (2014).

31. Li, H. et al. The landscape of cancer cell line metabolism. Nat. Med. 25, 850–860 (2019).

32. Ghandi, M. et al. Next-generation characterization of the Cancer Cell Line Encyclopedia. Nature 569, 503–508 (2019).

33. Fathi, A. T. et al. Isocitrate dehydrogenase 1 (IDH1) mutation in breast adenocarcinoma is associated with elevated levels of serum and urine 2-hydroxyglutarate. Oncologist 19, 602–607 (2014).

34. Chiang, S. et al. IDH2 mutations define a unique subtype of breast cancer with altered nuclear polarity. Cancer Res. 76, 7118–7129 (2016).

35. Bhargava, R. et al. Breast tumor resembling tall cell variant of papillary thyroid carcinoma: A solid papillary peoplasm with characteristic immunohistochemical profile and few recurrent mutations. Am. J. Clin. Pathol. 147, 399–410 (2017).

36. Branco, M. R., Ficz, G. & Reik, W. Uncovering the role of 5-hydroxymethylcytosine in the epigenome. Nat. Rev. Genet. 13, 7–13 (2011).

37. Buenrostro, J. D. et al. Single-cell chromatin accessibility reveals principles of regulatory variation. Nature 523, 486–490 (2015).

38. Cusanovich, D. A. et al. Multiplex single cell profiling of chromatin accessibility by combinatorial cellular indexing. Science 348, 910–914 (2015).

39. Davis, C. A. et al. The Encyclopedia of DNA elements (ENCODE): data portal update. Nucleic Acids Res. 46, D794–d801 (2018).

40. Ernst, J. et al. Mapping and analysis of chromatin state dynamics in nine human cell types. Nature 473, 43–49 (2011).

41. Gu, F. et al. CMS: a web-based system for visualization and analysis of genome-wide methylation data of human cancers. PLoS One 8, e60980 (2013).

42. Cancer Genome Atlas, N. Comprehensive molecular portraits of human breast tumours. Nature 490, 61–70 (2012).

43. Ernst, J. & Kellis, M. Chromatin-state discovery and genome annotation with ChromHMM. Nat. Protoc. 12, 2478–2492 (2017).

44. Huang, W. Y. et al. MethHC: a database of DNA methylation and gene expression in human cancer. Nucleic Acids Res. 43, D856–861 (2015).

45. Kramer, A., Green, J., Pollard, J., Jr. & Tugendreich, S. Causal analysis approaches in Ingenuity Pathway Analysis. Bioinformatics 30, 523–530 (2014).

46. Corces, M. R. et al. The chromatin accessibility landscape of primary human cancers. Science 362 (2018).

47. Adam, M., Robert, F., Larochelle, M. & Gaudreau, L. H2A.Z is required for global chromatin integrity and for recruitment of RNA polymerase II under specific conditions. Mol. Cell. Biol. 21, 6270–6279 (2001).

48. Jadhav, R. R. et al. Genome-wide DNA methylation analysis reveals estrogen-mediated epigenetic repression of metallothionein-1 gene cluster in breast cancer. Clin. Epigenetics 7, 13 (2015).

49. Bendall, S. C. et al. Single-cell mass cytometry of differential immune and drug responses across a human hematopoietic continuum. Science 332, 687–696 (2011).

50. Cheung, P. et al. Single-cell chromatin modification profiling reveals increased epigenetic variations with aging. Cell 173, 1385–1397.e1314 (2018).

51. Lord, C. J. & Ashworth, A. BRCAness revisited. Nat. Rev. Cancer 16, 110–120 (2016).

52. Wang, P. et al. Oncometabolite D-2-hydroxyglutarate inhibits ALKBH DNA repair enzymes and sensitizes IDH mutant cells to alkylating agents. Cell Rep. 13, 2353–2361 (2015).

53. Chen, F. et al. Oncometabolites d- and l-2-hydroxyglutarate inhibit the AlkB family DNA repair enzymes under physiological conditions. Chem. Res. Toxicol. 30, 1102–1110 (2017).

54. Inoue, S. et al. Mutant IDH1 downregulates ATM and alters DNA repair and sensitivity to DNA damage independent of TET2. Cancer Cell 30, 337–348 (2016).

55. Sulkowski, P. L. et al. 2-Hydroxyglutarate produced by neomorphic IDH mutations suppresses homologous recombination and induces PARP inhibitor sensitivity. Sci. Transl. Med. 9 (2017).

56. Flavahan, W. A. et al. Insulator dysfunction and oncogene activation in IDH mutant gliomas. Nature 529, 110–114 (2016).

57. Yan, H. et al. IDH1 and IDH2 mutations in gliomas. N. Engl. J. Med. 360, 765–773 (2009).

58. Hartmann, C. et al. Long-term survival in primary glioblastoma with versus without isocitrate dehydrogenase mutations. Clin. Cancer Res. 19, 5146–5157 (2013).

59. Chakraborty, A. A. et al. Histone demethylase KDM6A directly senses oxygen to control chromatin and cell fate. Science 363, 1217–1222 (2019).

60. Batie, M. et al. Hypoxia induces rapid changes to histone methylation and reprograms chromatin. Science 363, 1222–1226 (2019).

61. Neve, R. M. et al. A collection of breast cancer cell lines for the study of functionally distinct cancer subtypes. Cancer Cell 10, 515–527 (2006).

62. Hsu, P. Y. et al. Amplification of distant estrogen response elements deregulates target genes associated with tamoxifen resistance in breast cancer. Cancer Cell 24, 197–212 (2013).

63. Lin, A. P. et al. D2HGDH regulates alpha-ketoglutarate levels and dioxygenase function by modulating IDH2. Nat. Commun. 6, 7768 (2015).

64. Struys, E. A., Jansen, E. E., Verhoeven, N. M. & Jakobs, C. Measurement of urinary D- and L-2-hydroxyglutarate enantiomers by stable-isotope-dilution liquid chromatography-tandem mass spectrometry after derivatization with diacetyl-L-tartaric anhydride. Clin. Chem. 50, 1391–1395 (2004).

65. Yu, M. et al. Tet-assisted bisulfite sequencing of 5-hydroxymethylcytosine. Nat. Protoc. 7, 2159–2170 (2012).

66. Mitsuya, K. et al. Alterations in the placental methylome with maternal obesity and evidence for metabolic regulation. PLoS One 12, e0186115 (2017).

67. Garcia, B. A. et al. Chemical derivatization of histones for facilitated analysis by mass spectrometry. Nat. Protoc. 2, 933–938 (2007).

68. Camarillo, J. M. et al. Coupling fluorescence-activated cell sorting and targeted analysis of histone modification profiles in primary human leukocytes. J. Am. Soc. Mass Spectrom. (2019).

69. Diebold, L. P. et al. Mitochondrial complex III is necessary for endothelial cell proliferation during angiogenesis. Nat. Metab. 1, 158–171 (2019).

70. Langmead, B. & Salzberg, S. L. Fast gapped-read alignment with Bowtie 2. Nat. Methods 9, 357–359 (2012).

71. Ji, Z., Zhou, W. & Ji, H. Single-cell regulome data analysis by SCRAT. Bioinformatics 33, 2930–2932 (2017).

72. Consortium, E. P. An integrated encyclopedia of DNA elements in the human genome. Nature 489, 57–74 (2012).

73. Gao, J. et al. Integrative analysis of complex cancer genomics and clinical profiles using the cBioPortal. Sci. Signal. 6, pl1 (2013).

74. Booth, M. J. et al. Quantitative sequencing of 5-methylcytosine and 5-hydroxymethylcytosine at single-base resolution. Science 336, 934–937 (2012).

75. Takahashi, C. et al. Mass cytometry panel optimization through the designed distribution of signal interference. Cytometry A 91, 39–47 (2017).

76. Hartmann, F. J. et al. Scalable conjugation and characterization of Immunoglobulins with stable mass isotope reporters for single-cell mass cytometry analysis. Methods Mol. Biol. 1989, 55–81 (2019).

77. Kotecha, N., Krutzik, P. O. & Irish, J. M. Web-based analysis and publication of flow cytometry experiments. Curr. Protoc. Cytom. Chapter 10, Unit10.17 (2010).

78. Finck, R. et al. Normalization of mass cytometry data with bead standards. Cytometry A 83, 483–494 (2013).

