## Supplementary Figures for "Single-cell chromatin profiling reveals demethylation-dependent metabolic vulnerabilities of breast cancer epigenome"

Supplementary Figure 1

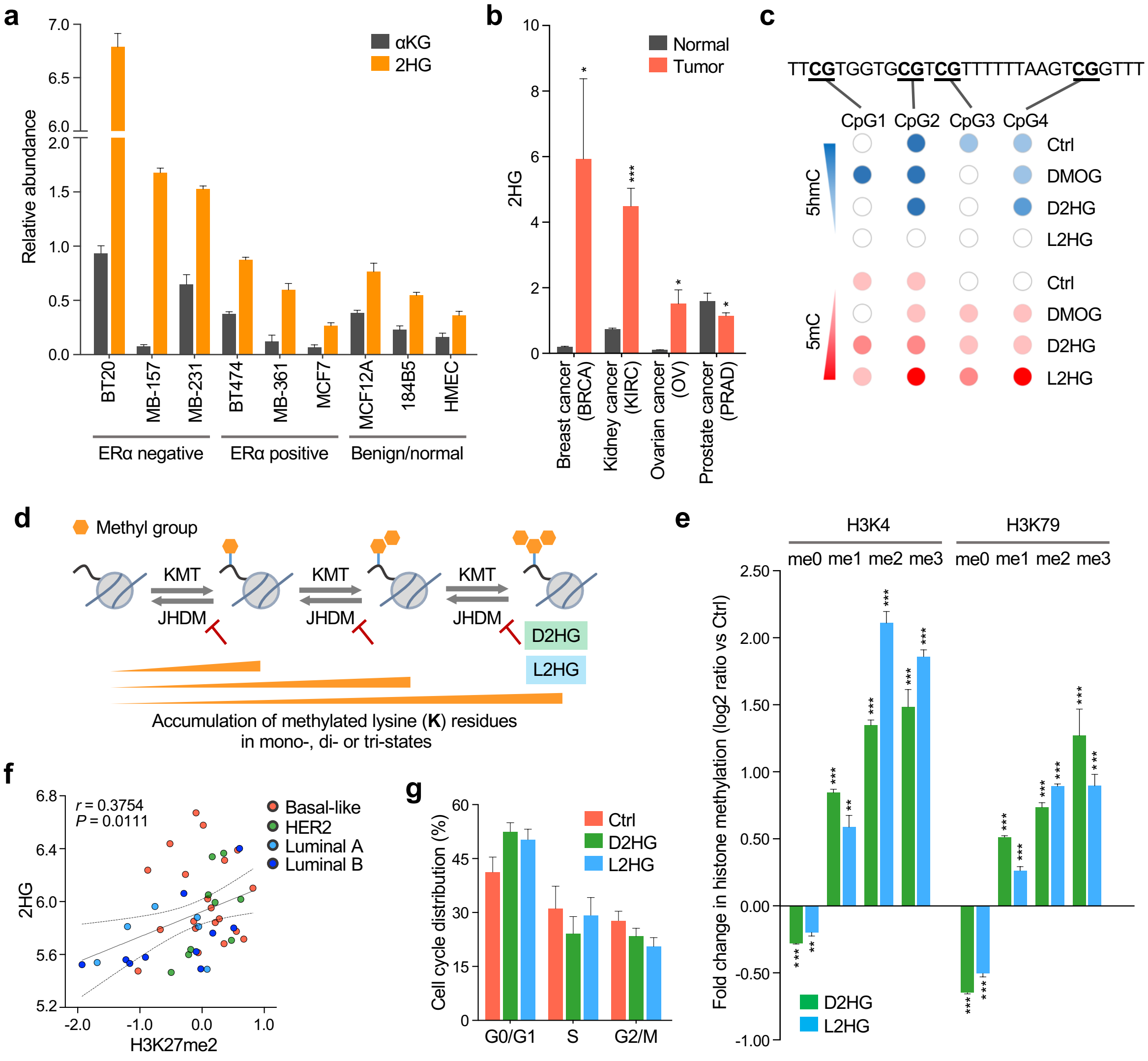

**Supplementary Figure 1. Intratumor 2HG progressively modulates the mammary epithelial epigenome independent of cell cycle. a**, LC-MS analysis of intracellular levels of αKG and 2HG in breast cancer cell lines, benign and normal mammary epithelial cells (n = 3 independent replicates). **b**, Relative abundance of 2HG in tumor and normal tissues across different cancer types. *$P < 0.05$, ***$P < 0.001$ by two-tailed unpaired Student's *t*-test: breast invasive carcinoma (BRCA), tumor (n = 92) vs. normal (n= 70); kidney renal clear cell carcinoma (KIRC), tumor (n = 138) vs. normal (n= 138); ovarian serous cystadenocarcinoma (OV), tumor (n = 18) vs. normal (n= 12); prostate adenocarcinoma (PRDA), tumor (n = 60) vs. normal (n= 25). In some, but not all cases, matched adjacent normal tissue data were available. **c**, 5hmC and 5mC levels at LINE-1 repetitive genomic sequences. HMECs were exposed for 72 hr to either 2 mM DMOG, 1 mM D2HG or L2HG, and subjected to TAB pyrosequencing in order to differentiate 5hmC from 5mC (see Methods for further details). DMOG is an analog of αKG and competes for binding at the active center of the enzyme. **d**, Schematic showing the accumulation of methylated lysine (K) residues by 2HG-mediated inhibition of histone demethylation. Histone lysine residues are mono-, di- and trimethylated by KMT (histone methyltransferases) and αKG-dependent JHDM (Jumonji-domain containing histone demethylases) remove methyl groups from the lysines. **e**, Fold change in different classes of histone methylation upon 2HG exposure. Following 72-hr exposure to 100 µM of either D2HG or L2HG, global levels of methylated/unmethylated lysine residues were assayed by multiplexed mass spectrometry. *$P < 0.05$, **$P < 0.01$, ***$P < 0.001$ versus unexposed control HMECs by two-tailed unpaired Student's *t*-test with Holm-Sidak correction for multiple comparisons (n = 3 independent replicates). **f**, Relationship between intratumor 2HG and H3K27me2 levels in the CCLE breast cancer cell lines. Solid and dashed lines represent linear fit and 95% confidence interval of the fitting respectively. Spearman's correlation coefficient (*r*) and corresponding *P* value are given at the top of the panel: basal-like (n = 20), HER2-enriched (n = 9), luminal A (n = 6) and luminal B (n = 10) breast tumor molecular subtypes (see Methods for details regarding the CCLE datasets). **g**, Cell cycle analysis of HMECs exposed for 72 hr to 100 µM of either D2HG or L2HG. No statistical significance was observed by one-way ANOVA with Dunnett's multiple comparison test. Results are from three independent experiments. Error bars in column charts denote s.e.m. LINE-1, long interspersed nuclear element type 1; DMOG, dimethyloxalylglycine; TAB, Tet-assisted bisulfite; CCLE, Cancer Cell Line Encyclopedia; me0, unmethylated; me1, monomethylated; me2, dimethylated; me3, trimethylated histone lysine residues; HMECs, human mammary epithelial cells; HER2, human epidermal growth factor receptor 2.

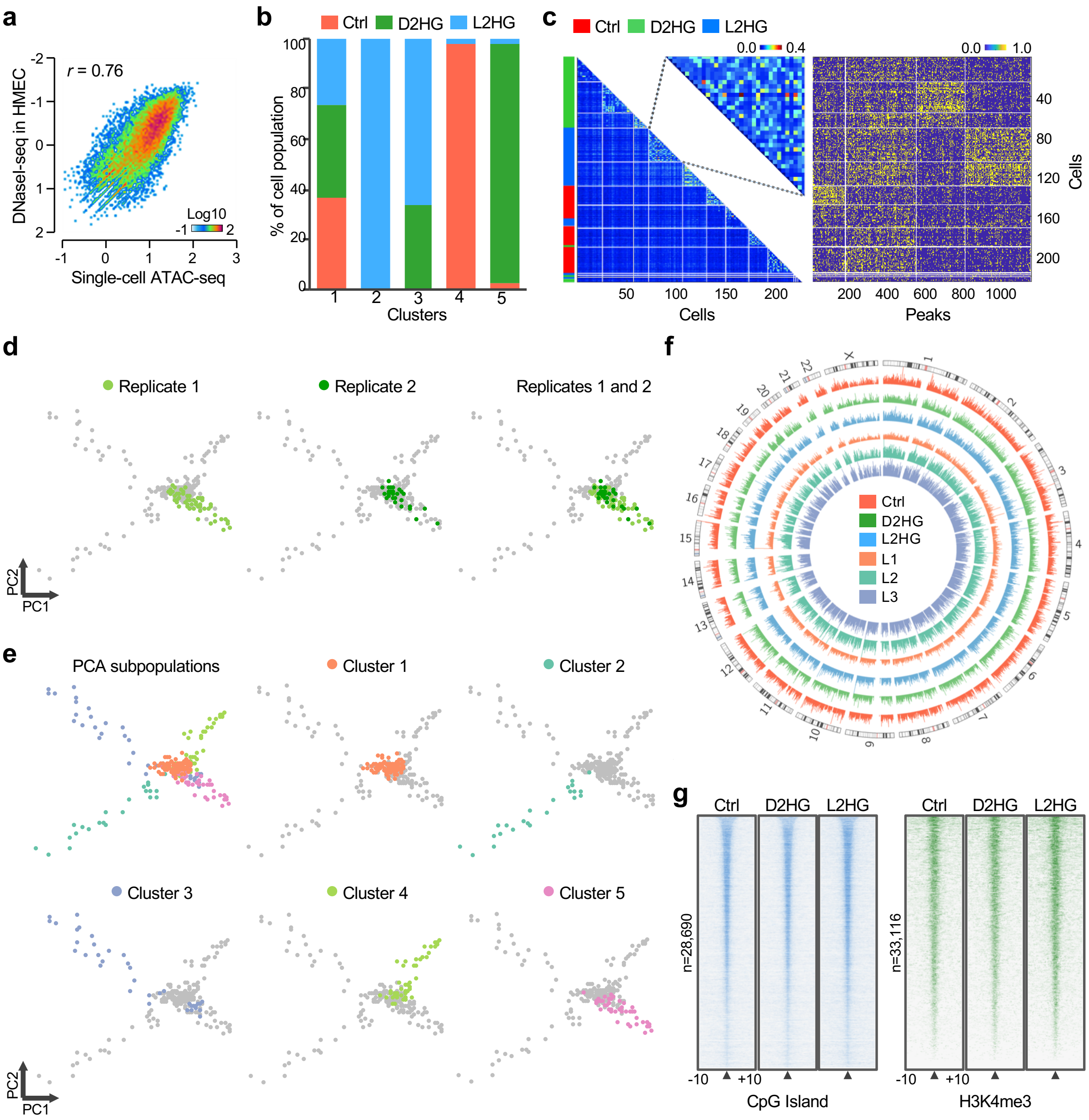
a
r = 0.76
DNaseI-seq in HMEC
Single-cell ATAC-seq
Log10
b
Ctrl D2HG L2HG
% of cell population
Clusters
c
Ctrl D2HG L2HG
Cells
Peaks
d
Replicate 1
Replicate 2
Replicates 1 and 2
PC2
PC1
e
PCA subpopulations
Cluster 1
Cluster 2
Cluster 3
Cluster 4
Cluster 5
f
Ctrl
D2HG
L2HG
L1
L2
L3
g
Ctrl D2HG L2HG
n=28,690
n=33,116
CpG Island
H3K4me3

**Supplementary Figure 2. 2HG enantiomers differentially modulate chromatin regulatory landscape of the mammary epithelial epigenome. a**, Comparison of aggregate scATAC-seq and ensemble DNaseI-seq signals. Genome-wide chromatin accessibility patterns observed by scATAC-seq are highly correlated with DNaseI-seq data (GSE29692) from HMECs (*r* = 0.76). **b**, Distribution of cell populations in different clusters identified by model-based clustering, colored by sample type. **c**, Epigenetic heterogeneity revealed by cell-to-cell similarity matrix (left) and corresponding open chromatin peaks (right) based on weighted associations between scATAC-seq fragments and commonly defined chromatin accessibility peaks. Network-based clustering was applied to ~1200 highly accessible peaks. **d**, A two-dimensional projection of the scATAC-data using PCA. Colors represent two different biological replicates, showing a high degree of consistency between batches. **e**, PCA projections of five distinct cell clusters identified by model-based clustering. Each data point represents a single cell. Cell clusters are color-coded as indicated. **f**, Circos plot depicting the genome-wide alterations in chromatin accessibility peaks in D2HG- and L2HG-exposed cells or among L2HG subpopulations. The outer three tracks represent control (red), D2HG-exposed (green) and L2HG-exposed (blue) cells, while the inner three tracks show chromatin accessibility peaks in L1 (orange), L2 (light green) and L3 (purple) subpopulations. Peak heights correspond to the degree of DNA accessibility. The outermost ring represents chromosome annotations. **g**, Whole-genome heat maps showing enrichment of scATAC-seq signals over a 20-kb region centered on CpG islands and H3K4me3 peak summits, sorted by decreasing scATAC-seq signal. Each row represents one individual genomic locus and the color represents the intensity of chromatin accessibility. Intervals flanking indicated feature are shown in kilobases. ATAC, assay for transposase accessible chromatin; PCA, principal component analysis.

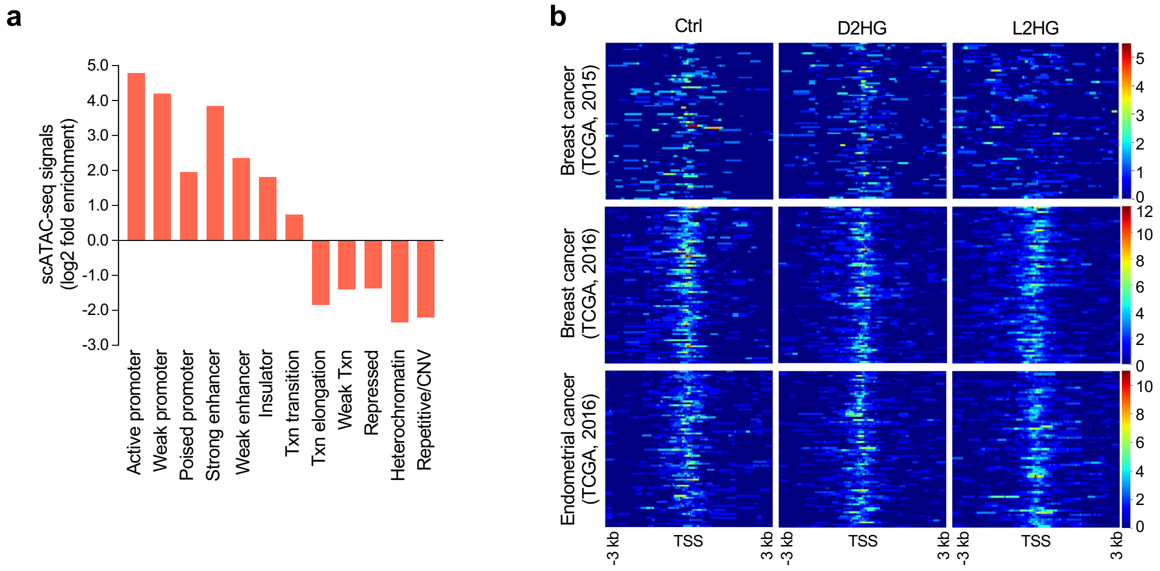


**Supplementary Figure 3. 2HG mediated chromatin remodeling is accompanied by loss of promoter accessibility.** **a**, Enrichment of scATAC-seq signals in ChromHMM-defined chromatin states relative to values expected by chance. **b**, Heat maps showing the chromatin accessibility at gene promoters that are highly methylated in the TCGA cancer cohorts. scATAC-seq profiles in 2HG-exposed and unexposed cells are centered on TSS of highly methylated genes (n = 150 genes) in breast cancer (two independent datasets) and endometrial cancer patients. Each row represents a single cell. ChromHMM, chromatin Hidden Markov Modeling; TCGA, The Cancer Genome Atlas; TSS, transcriptional start site.

### Supplementary Figure 4

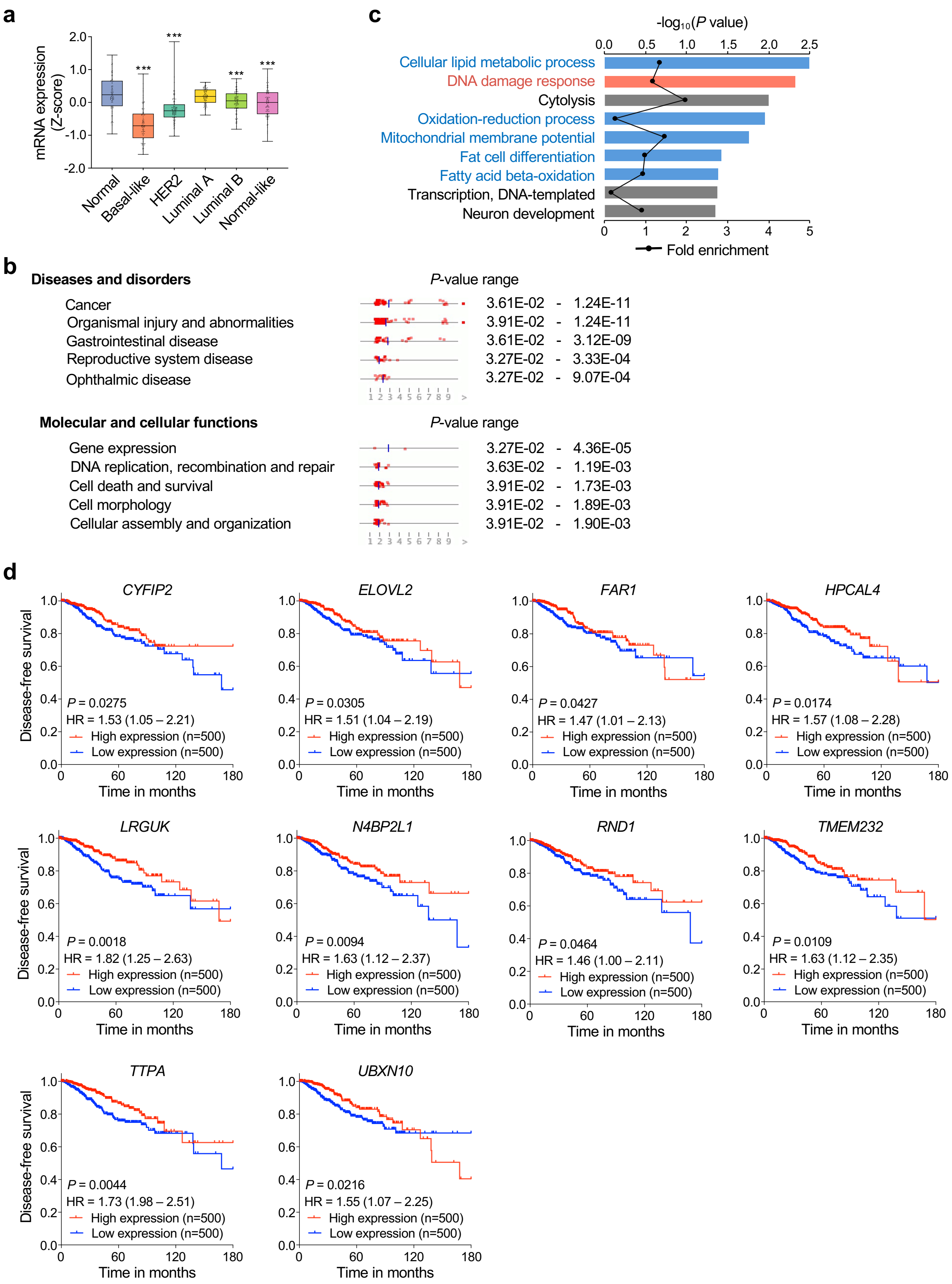

**Supplementary Figure 4. Tumor hypermethylation is associated with downregulation of genes involved in cellular metabolic processes and DNA damage responses.** **a**, Box and whiskers plot with center line representing median mRNA expression of genes that are highly methylated (n = 150) in the TCGA breast cancer dataset. Boxplot boundaries indicate the first and third quartiles of the data points and whiskers extend to the furthest data point. ***$P < 0.001$ versus adjacent normal tissue by one-way ANOVA with Dunnett's multiple comparison test. Adjacent normal tissue (n = 112), basal-like (n = 97), HER2-enriched (n = 58), luminal A (n = 231), luminal B (n = 127) and normal-like (n = 8) subtypes (see Methods for further details). **b**, Enriched disease and biofunctional pathways identified by IPA. Most significant categories are shown with associated *P* values. **c**, GO biological processes identified by enrichment analysis. DAVID functional annotation clustering tool (http://david.abcc.ncifcrf.gov) was used to detect significantly overrepresented GO biological processes. **d**, Kaplan-Meier analysis of disease-free survival (DFS) in the TCGA breast cancer patients, stratified by mRNA expression of hypermethylated genes. The median expression value was used as a threshold to define high versus low expression. Log-rank (Mantel-Cox) *P* values and hazard ratios (HR) are shown. GO, Gene Ontology; DAVID, Database for annotation, visualization and integrated discovery; IPA, Ingenuity Pathways Analysis; TCGA, The Cancer Genome Atlas.

Supplementary Figure 5

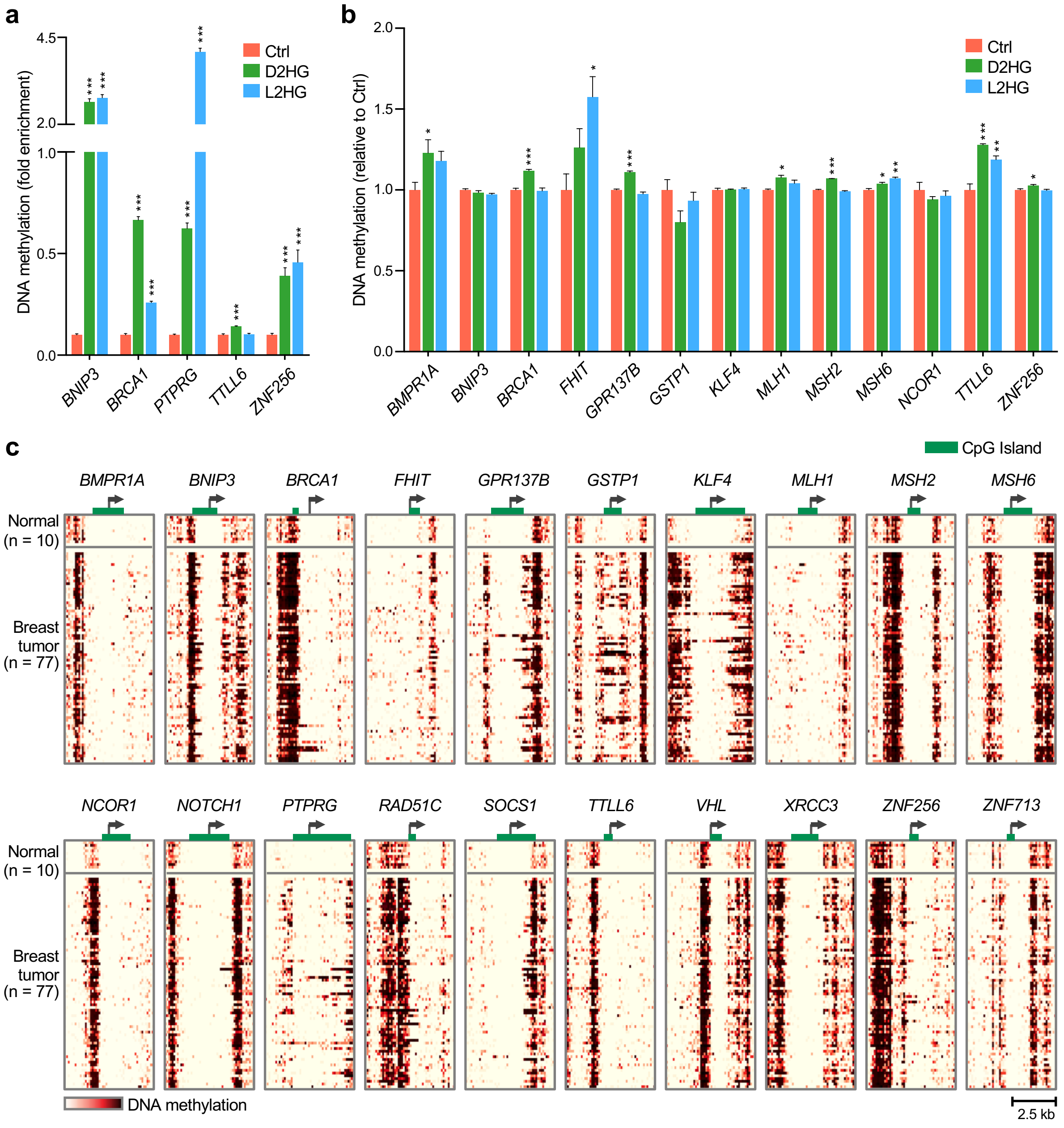

**Supplementary Figure 5. Promoter hypermethylation of tumor suppressor genes is induced upon 2HG exposure.** **a**, Promoter hypermethylation in 2HG-perturbed HMECs assessed by MeDIP-qPCR. Methylated DNA fragments were enriched using 5mC-specific antibody to differentiate 5mC from 5hmC. Results are normalized to input signals and values are expressed as fold enrichment relative to unexposed control cells. **b**, Promoter methylation in hTERT-HME1 cells measured by oxBS pyrosequencing. *$P < 0.05$, **$P < 0.01$, ***$P < 0.001$ versus control by one-way ANOVA with Dunnett's multiple comparison test ($n \geq 3$ individual replicates). Error bars represent s.e.m. **c**, Heat maps showing hypermethylated gene promoters in breast cancer. DNA methylation landscape was generated by MBDCap-seq using primary breast tumors (n = 77) and uninvolved breast tissue (n = 10). Green bars indicate locations of promoter CpG islands and arrows indicate TSS and the direction of transcription. MeDIP, methylated DNA immunoprecipitation; qPCR, quantitative polymerase chain reaction; oxBS, oxidative bisulfite; MBDCap-seq, methyl-binding domain capture sequencing; TSS, transcriptional start site.

Supplementary Figure 6

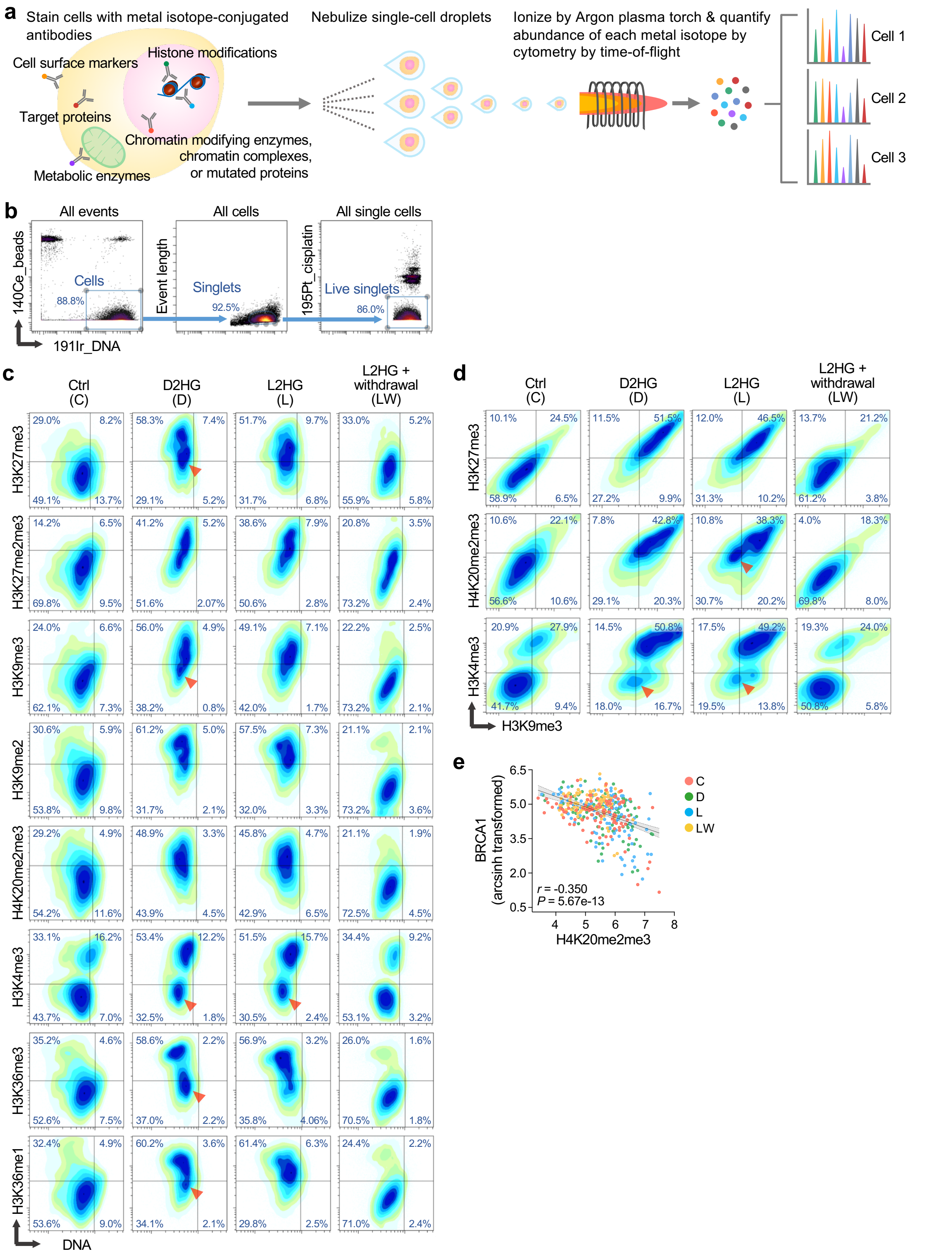

**Supplementary Figure 6. Mass cytometry demonstrates concurrent alterations in multiple distinct types of chromatin modifications and differentiates cell subpopulations with different sensitivity to 2HG exposure. a**, Workflow summary of single-cell epigenetic profiling by mass cytometry. Cells are stained with epitope-specific antibodies conjugated with different metal isotopes, nebulized into single-cell droplets and passed through an argon plasma where cells are vaporized, atomized and ionized to form clouds of ions that correspond to individual cells. Metal-labeled antibodies are used to simultaneously detect epigenetic modifications as well as cellular target proteins. **b**, Sequential gating strategy to isolate live intact single cells. A three-step Boolean gating approach was applied simultaneously to all samples. 191Ir and 140Ce were used to identify cells. Events were then gated on singlets according to DNA content and event length to exclude debris and cell aggregates. Subsequently, live single cells were gated based on cisplatin (195Pt) exclusion for downstream analysis. After gating, individual channels were visualized in biaxial plots. **c**, 2D contour plots showing an increase in methylated lysine residues following 2HG exposure and subsequent restoration of histone methylation upon 2HG withdrawal. C, unexposed control cells; D, cells exposed to D2HG for 72 hr; L, cells exposed to L2HG for 72 hr; LW, cells exposed to L2HG for 72 hr followed by 5-day withdrawal of L2HG. Arrowheads indicate cell populations that are potentially less sensitive to 2HG oncometabolite perturbation. **d**, 2D contour plots showing a concurrent increase in different types of histone methylation modifications following 2HG exposure. Arrowhead indicates cell populations that are potentially less sensitive to 2HG oncometabolite perturbation. Percentage of cells in each quadrant is indicated. **e**, Correlation of BRCA1 expression to H4K20 methylation levels. Each data point represents an individual cell in Cluster 1 and is colored according to the experimental groups. Data are transformed to arcsinh scales with the cofactor of 5. Solid and shaded area represent linear fit and 95% confidence interval of the fitting respectively. Spearman's correlation coefficient (*r*) and corresponding *P* value are given at the bottom left. 2D, two-dimensional; HMECs, human mammary epithelial cells.
